# Nasopharyngeal competition dynamics are likely to be altered following vaccine introduction: bacteriocin prevalence and diversity among Icelandic and Kenyan pneumococci

**DOI:** 10.1101/2022.11.03.514821

**Authors:** Madeleine EB Butler, Melissa J Jansen van Rensburg, Angela Karani, Benedict Mvera, Donald Akech, Asma Akter, Calum Forrest, Andries J van Tonder, Sigríður J Quirk, Gunnsteinn Haraldsson, Stephen D Bentley, Helga Erlendsdóttir, Ásgeir Haraldsson, Karl G Kristinsson, J Anthony G Scott, Angela B Brueggemann

**Affiliations:** Imperial College London, London, United Kingdom; University of Oxford, Oxford, United Kingdom; KEMRI Wellcome Trust Programme, Kilifi, Kenya; University of Cambridge, Cambridge, United Kingdom; University of Iceland and Landspitali - The National University Hospital of Iceland, Reykjavík, Iceland; Wellcome Sanger Institute, Hinxton, United Kingdom; University of Iceland and Children’s Hospital Iceland, Reykjavík, Iceland; London School of Hygiene and Tropical Medicine, London, United Kingdom

## Abstract

Bacteriocins are antimicrobial peptides produced by bacteria to inhibit other bacteria in the surrounding environment. *Streptococcus pneumoniae* is a leading cause of disease worldwide and colonises the healthy human nasopharynx, where it competes for space and nutrients. Pneumococcal conjugate vaccines have reduced the incidence of disease, but they also restructure the bacterial population, and this restructuring likely alters the nasopharyngeal competition dynamics. Here, the distribution of bacteriocins was examined in over 5000 carriage and disease-causing pneumococci from Iceland and Kenya, recovered before and after the introduction of pneumococcal vaccination. Overall, up to eleven different bacteriocin gene clusters were identified per pneumococcus. Significant differences in the prevalence of bacteriocins were observed before and after vaccine introduction, and among carriage and disease-causing pneumococci, which were largely explained by the bacterial population structure. Genetically similar pneumococci generally harboured the same bacteriocins although sometimes different repertoires of bacteriocins were observed, which suggested that horizontal transfer of bacteriocin clusters had occurred. These findings demonstrated that vaccine-mediated changes in the pneumococcal population altered the prevalence and distribution of bacteriocins. The consequences of this for pneumococcal colonisation and disease remain to be determined.

## Introduction

*Streptococcus pneumoniae* (pneumococcus), is a major cause of lower respiratory tract infections (LRTI), invasive pneumococcal disease (IPD) including meningitis and bacteraemia, and otitis media (OM). Despite vaccines and antimicrobials, in 2016 it was estimated that pneumococcus caused 197 million episodes of pneumonia and more than 1.1 million deaths worldwide, disproportionately affecting low- and middle-income countries^1^.

The ecological niche of the pneumococcus is the paediatric nasopharynx and whilst carriage is usually asymptomatic it is a precursor to disease^2^. Carriage duration varies between serotypes and less invasive serotypes typically colonise for longer periods of time^3^. Since pneumococci and other bacterial species in the nasopharynx are competing for space and resources, competition dynamics are likely to be associated with pneumococcal pathogenesis, although the mechanism of competition is not yet understood^4,5^. Furthermore, pneumococcal conjugate vaccines (PCVs) target a limited number of serotypes, and after PCV implementation vaccine serotypes typically decrease and nonvaccine serotypes increase within the human population, which results in changes to nasopharyngeal competition dynamics^6-8^.

Bacteria can compete with bacteriocins, which are peptides or small proteins that give the producing bacteria an advantage by killing or inhibiting other bacteria in the environment^9-11^. Bacteriocins are encoded on biosynthetic gene clusters, which in the simplest form encode a toxin plus an ‘immunity’ gene(s) to protect the producing strain from its own bacteriocin. Bacteriocin clusters may also encode genes for post-translational modification enzymes, dedicated transporters, or regulatory systems^11^. The most well-studied pneumococcal bacteriocin is the bacteriocin-like peptide (*blp*) cluster, which is highly diverse and ubiquitous.^9,12,13^. More recently, genome mining studies revealed a wide repertoire of different bacteriocin clusters among large and diverse pneumococcal datasets; however, the prevalence and diversity of bacteriocins within well-defined subsets of the pneumococcal population has not been investigated.^12,14,15^

Therefore, the aim of this study was to investigate bacteriocin clusters among pneumococci recovered from disease and carriage in Iceland and Kenya, before and after the introduction of 10-valent PCV (PCV10) immunisation. In total, over 5000 pneumococcal genomes were screened for 20 different bacteriocin clusters, which revealed that bacteriocins were widespread, were associated with the underlying pneumococcal population structure in each country, and that the prevalence and distribution of these bacteriocins were significantly altered post-PCV10. Overall, these findings suggest that nasopharyngeal competition dynamics are likely to be altered following PCV introduction.

## Methods

### Pneumococcal isolate collections, microbiology, and serotyping

Icelandic carriage pneumococci were recovered from nasopharyngeal swabs of healthy children <7 years of age in day care centres in the Reykjavik area. Swabs were collected in March every year from 2009 to 2014, except in 2009 when 40% of the samples were taken in April. Disease pneumococci were recovered from diagnostic specimens sent to the clinical microbiology laboratory at Landspitali University Hospital from 2009 to 2014: OM isolates were collected from the middle ear of children <7 years of age with OM; LRTI isolates were collected from the sputum of adults (>18 years of age) with suspected pneumonia; and invasive pneumococci were collected from normally sterile sites of patients of all ages with IPD. All of the Icelandic IPD isolates collected between 2009 and 2014, and every other carriage, LRTI and OM isolate in the overall dataset were selected for whole genome sequencing. The pneumococci recovered from carriage, LRTI and OM were published previously^7,8^.

Kenyan isolates were selected for whole genome sequencing from pneumococcal collections at the KEMRI-Wellcome Trust Research Programme, and were recovered from residents of the Kilifi Health and Demographic Surveillance System (KHDSS)^6^. Carriage pneumococci were recovered from healthy patients of all ages in the KHDSS who had been recruited to carriage studies in the district^6,16-19^. Isolates collected between 2004 and 2017, excluding 2011 (the year PCV10 was introduced), were sampled randomly from the age groups represented by the carriage studies: 3-5, 6-11, 12-23 and 24-50 months, 5-14, 15-64 and 65+ years. Sampling within each age stratum was weighted to reflect the observed population structure of residents of the KHDSS. Invasive pneumococci were collected from patients of all ages presenting with IPD at Kilifi County Hospital, and all isolates collected from 2003 to 2017 were included in this study.

All pneumococci were cultured from primary specimens and identified using standard microbiological techniques. Isolates were serotyped in Kenya using latex agglutination and confirmed with the Quellung reaction if necessary. Pneumococci in Iceland were serotyped using latex agglutination and confirmed with multiplex PCR assays and sequence-based serotyping when needed^7,8,20,21^

### DNA extraction and whole genome sequencing

Freezer stocks of pneumococcal isolates were cultured using standard microbiological methods. Optochin disk susceptibility confirmed that isolates were pneumococci and any viridans streptococci were excluded from further processing. DNA was extracted as described previously^7^ and sequenced using the Illumina HiSeq2000 platform.

### Genome assembly, quality control, MLST and clonal complex definitions

Draft genomes were assembled *de novo* into contigs from short paired-end reads using an internal Sanger pipeline that utilised Velvet Optimiser, SSPACE and GapFiller^22-24^. Isolate records with corresponding provenance data and assembled genomes were stored in a private PubMLST database for analysis during this study^25^. Sequence assembly statistics (ie genome size, number of contigs, GC content) were inspected and any genome with a value greater than two standard deviations from the mean of the dataset for any of the assembly statistics was investigated manually.

The ribosomal multilocus sequence typing (rMLST) species identification tool (https://pubmlst.org/species-id) was used to confirm that genomes were pneumococcal sequences^26^. Within PubMLST, genomes were automatically screened and assigned multilocus sequence typing (MLST) and rMLST alleles, and any new alleles and sequence types (STs) were submitted to PubMLST curators for assignment (https://pubmlst.org/organisms/streptococcus-pneumoniae). MLST and rMLST data were investigated manually to identify any locus with >1 allele assignment, which suggested contamination. Suboptimal genomes were excluded from further analyses.

Phyloviz was used to define clonal complexes (CCs) based upon founder STs and closely related single locus variants (SLVs), using the entire PubMLST database of unique STs. CCs were named after the founder ST^27^. A pneumococcus designated a ‘singleton’ did not have any SLVs that connected it to any other ST.

### *In silico* serotyping

Icelandic pneumococci were previously serotyped *in silico* with the seqSerotyper tool^7,20^ and SeroBA was used to predict the serotypes of the Kenyan genomes^28^. Discrepancies between phenotypic and *in silico* serotypes in the Kenyan dataset were resolved manually by comparisons to reference capsular polysaccharide (*cps*) locus sequences, but if this was unclear then the phenotypic serotype was used in analyses.

### Annotation of bacteriocin coding sequences

Previously described bacteriocin coding sequences (hereafter simply referred to as ‘genes’) were defined within PubMLST and then study genomes were automatically screened for these genes using BLAST searches^25^. Alleles were manually curated, and assigned to full length bacteriocin gene sequences and those with insertions and deletions. A bacteriocin gene was considered to be absent if there was no match to an existing allele at >50% sequence identity and at least 30% gene length.

Streptolancidins A and J each contained a pair of small genes that were indistinguishable at a sequence level (*slaA1* and *slaA2, sljA1* and *sljA2*), and two genes from streptolancidin E (*sleT* and *sleX1*) were typically found as a small fragment elsewhere in the pneumococcal genome. The location of each set of genes was determined using the *in silico* hybridisation tool within PubMLST, and each gene was annotated based on its location within the genome. The streptolancidin E genes were annotated only when they occurred as part of a full streptolancidin E cluster.

### Validation of bacteriocin biosynthetic gene clusters

Full and partial bacteriocin biosynthetic gene clusters (hereafter ‘bacteriocin clusters’) were analysed: full clusters possessed the expected set of genes for that cluster; and partial clusters were incomplete but more than half of the expected genes were present. Partial clusters were included in analyses because previous experimental work demonstrated that genes in partial bacteriocin clusters were expressed^14^. Single bacteriocin cluster genes and bacteriocin cluster fragments that contained fewer than half of the expected genes were excluded from further analyses.

Because the bacteriocin alleles were assigned gene-by-gene, it was necessary to ensure that the bacteriocin genes were assembled as contiguous clusters. This was assessed computationally and a bacteriocin cluster was deemed contiguous if there were no gaps >2.5 Kb between each of the constituent bacteriocin cluster genes (Supplementary Figure 1). Bacteriocin clusters that did not meet these criteria were excluded from further analyses. Significant differences in bacteriocin cluster prevalence were determined using the Chi-square test. Figures were generated using Affinity Designer v.1.10.1 (https://affinity.serif.com/en-gb/designer/).

## Results

### Genomic datasets for bacteriocin analyses

Overall, 5,071 pneumococcal genomes were analysed for bacteriocin clusters. 1,912 Icelandic pneumococcal genomes were included: 1,733 carriage and non-invasive disease pneumococci that were previously published (although four were lower quality genome assemblies and excluded from these bacteriocin analyses)^7,8,20,21^; plus 183 new genomes of pneumococci recovered from patients with invasive disease and collected over the same time period (Table 1). In total, 3,159 Kenyan genomes were analysed: 3,372 pneumococci were selected for inclusion in this study; 3,258 genomes were sequenced; and 99 genomes were excluded following QC assessment (mainly duplicate genomes).

**Table 1:**
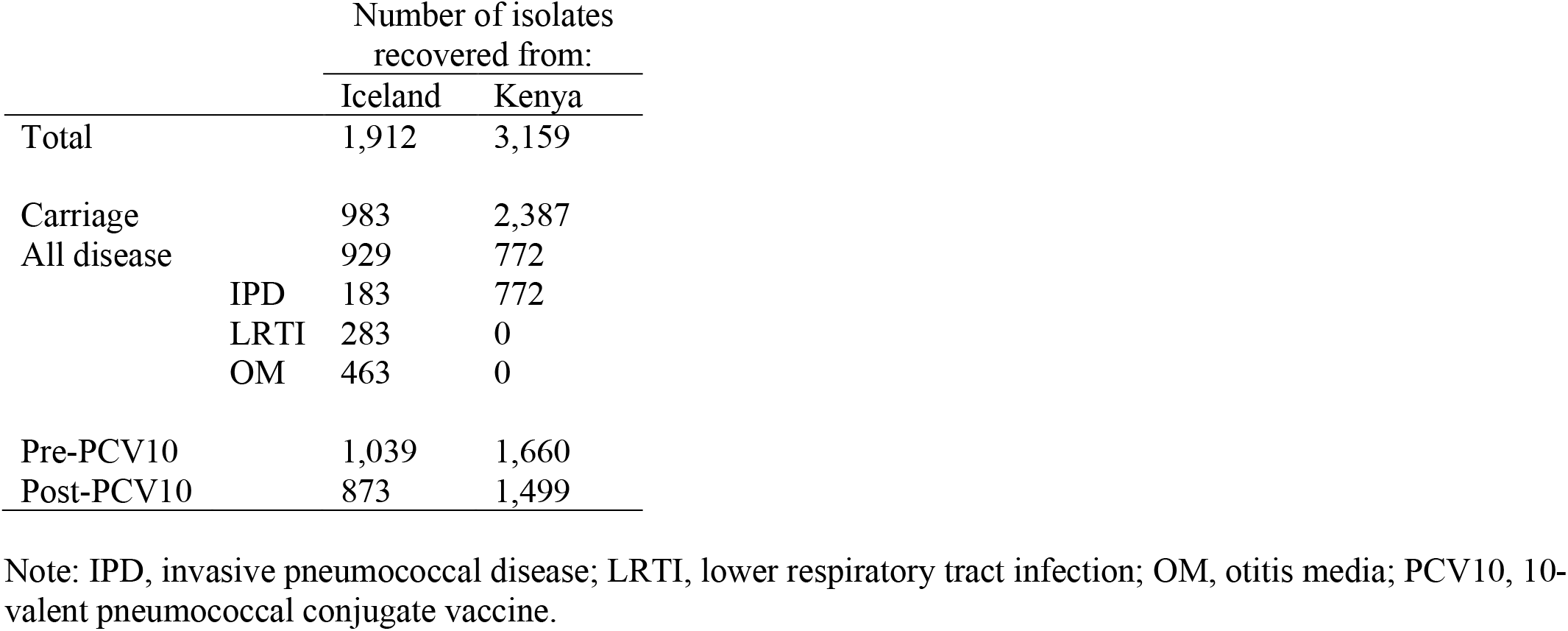
Pneumococcal genomes in the Icelandic and Kenyan study datasets.

Pneumococci were collected over six years (2009-2014) in Iceland and 14 years (2003-2017) in Kenya (Figure 1A). PCV10 was included in national infant immunisation programmes from 2011 onwards in both countries; therefore, the pre-vaccination period was the years up to and including 2011, and the post-vaccination period was from 2012 onwards. Disease-causing pneumococci were recovered from patients of all ages in both countries, whereas carriage pneumococci were recovered from children <7 years of age in Iceland, and from participants of all ages in Kenya (Figure 1B).

**Figure 1:**
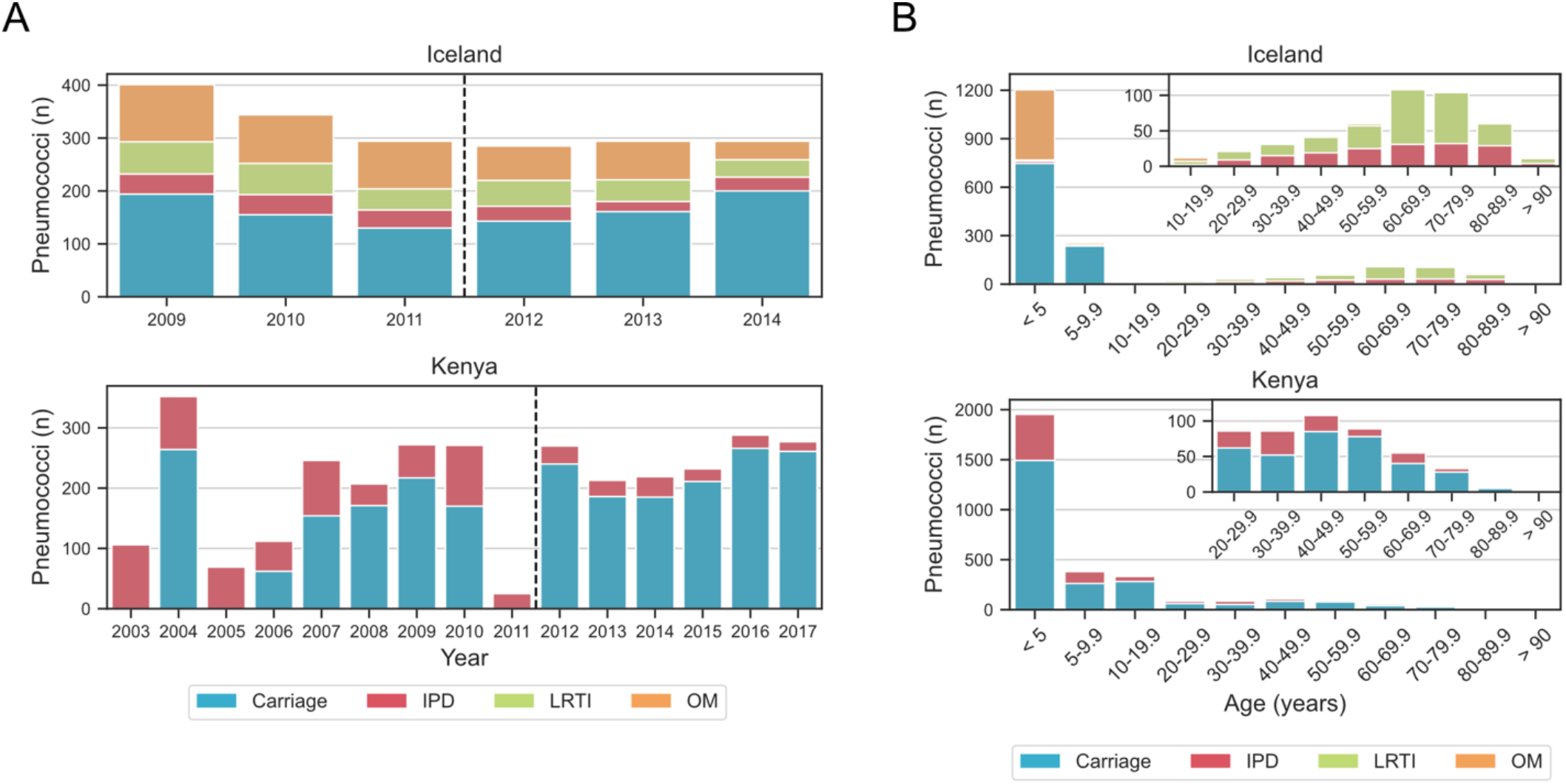
Description of the Icelandic and Kenyan datasets. (A) Numbers of carriage and disease pneumococci collected by year of isolation. The dashed line separates pre- and post-PCV10 periods. (B) Number of pneumococci collected by age group of study subjects. Inset plots increase data resolution for the older age groups. Note: IPD, invasive pneumococcal disease; LRTI, lower respiratory tract infection; OM, otitis media.

### Icelandic and Kenyan pneumococci represent distinct populations

59 unique CCs (and 5 singletons) were observed in the Icelandic dataset, and 116 unique CCs (and 62 singletons) were observed in the Kenyan dataset (Table 2). There were 18 CCs that were represented in both datasets. Overall, 40 and 56 serotypes were observed in the Icelandic and Kenyan datasets, respectively. The most prevalent serotype in both countries was 19F, which was dominant in the Icelandic dataset due to the incidence of OM caused by CC236/271/320 pneumococci in Iceland during the study period^7^. Other serotypes differed between the two populations, eg the highly invasive serotype 1 was nearly equal in prevalence to serotype 19F in the Kenyan dataset but serotype 1 was rare in the Icelandic dataset (Table 3).

**Table 2:** The 20 most prevalent clonal complexes (CCs) in the Icelandic and Kenyan datasets.

| **Iceland** |  | **Kenya** |  |
| --- | --- | --- | --- |
| **CC** | **n (%)** | **CC** | **n (%)** |
| 236/271/320 | 293 (15.3) | 5902 | 239 (7.6) |
| 439 | 217 (11.3) | 217 | 223 (7.1) |
| 199 | 179 (9.4) | 701 | 163 (5.2) |
| 138/176 | 122 (6.4) | 5339 | 142 (4.5) |
| 180 | 107 (5.6) | 1146 | 139 (4.4) |
| 62 | 94 (4.9) | 138/176 | 133 (4.2) |
| 97 | 87 (4.6) | 156/162 | 131 (4.1) |
| 490 | 74 (3.9) | 991 | 104 (3.3) |
| 124 | 62 (3.2) | 230 | 92 (2.9) |
| 433 | 61 (3.2) | 852 | 78 (2.5) |
| 30 | 60 (3.1) | 5258 | 77 (2.4) |
| 392 | 47 (2.5) | 63 | 70 (2.2) |
| 156/162 | 46 (2.4) | 289 | 69 (2.2) |
| 344 | 37 (1.9) | 347 | 62 (2.0) |
| 15 | 36 (1.9) | 914 | 61 (1.9) |
| 1262 | 35 (1.8) | 7053 | 58 (1.8) |
| 448 | 29 (1.5) | 702 | 58 (1.8) |
| 193 | 28 (1.5) | 854 | 57 (1.8) |
| 100 | 25 (1.3) | 499 | 55 (1.7) |
| 90 | 22 (1.2) | 1381 | 49 (1.6) |
| Other CCs<sup>a</sup> | 235 (12.3) | Other CCs<sup>a</sup> | 954 (30.2) |
| Singletons | 16 (0.8) | Singletons | 145 (4.6) |

a. 'Other CCs' represent 44 CCs in Iceland and 158 CCs in Kenya.

**Table 3:** The 20 most prevalent serotypes in the Icelandic and Kenyan datasets.

| **Iceland** |  | **Kenya** |  |
| --- | --- | --- | --- |
| **Serotype** | **n (%)** | **Serotype** | **n (%)** |
| 19F | 331 (17.3) | 19F | 228 (7.2) |
| 23F | 180 (9.4) | 1 | 224 (7.1) |
| 6A | 163 (8.5) | 6A | 206 (6.5) |
| 19A | 145 (7.6) | 19A | 153 (4.8) |
| 6B | 122 (6.4) | 15BC | 146 (4.6) |
| 3 | 109 (5.7) | 35B | 139 (4.4) |
| 11A | 93 (4.9) | 15A | 136 (4.3) |
| 15BC | 93 (4.9) | 14 | 131 (4.1) |
| 14 | 90 (4.7) | 6E(6Bii) | 131 (4.1) |
| nontypable | 70 (3.7) | 23F | 119 (3.8) |
| 22F | 61 (3.2) | 11A | 117 (3.7) |
| 23A | 51 (2.7) | 13 | 98 (3.1) |
| 23B | 47 (2.5) | 16F | 96 (3.0) |
| 35B | 33 (1.7) | 23B | 90 (2.8) |
| 9V | 32 (1.7) | 3 | 90 (2.8) |
| 21 | 29 (1.5) | 34 | 89 (2.8) |
| 6C | 29 (1.5) | 10A | 82 (2.6) |
| 33F | 29 (1.5) | 5 | 69 (2.2) |
| 6E(6Bii) | 26 (1.4) | 9V | 63 (2.0) |
| 16F | 26 (1.4) | 21 | 62 (2.0) |
| Other serotypes<sup>a</sup> | 153 (8.0) | Other serotypes | 690 (21.8) |

a. 'Other serotypes' represents additional serotypes in Iceland (n=20) and Kenya (n=36).

### Bacteriocin clusters were widely distributed among Icelandic and Kenyan pneumococci

In total, the presence or absence of 116 genes associated with 20 putative bacteriocin clusters was investigated among 5,071 pneumococcal genomes (Supplementary Table 3)^12,14,29-32^. Partial clusters were consistently observed for streptococcins B and E, and streptolancidins B, C, E and J (Supplementary Table 4). 95-100% of bacteriocin genes were in contiguous clusters (Supplementary Table 5).

Overall, cib and streptococcin B, C and E clusters were present in 96-100% of pneumococci from both countries, whereas streptolancidin H and I clusters were never observed (Figure 2A). The remaining bacteriocin clusters ranged in prevalence from 0.1% to 81% per dataset. Twelve bacteriocin clusters were significantly more common in one dataset than the other but the largest differences in prevalence were among the streptolancidins (Figure 2A). For example, 10.7% (n=340 genomes) of Kenyan pneumococci harboured a streptolancidin B cluster as compared to just two Icelandic pneumococci (Figure 2A; Supplementary Table 6).

**Figure 2:**
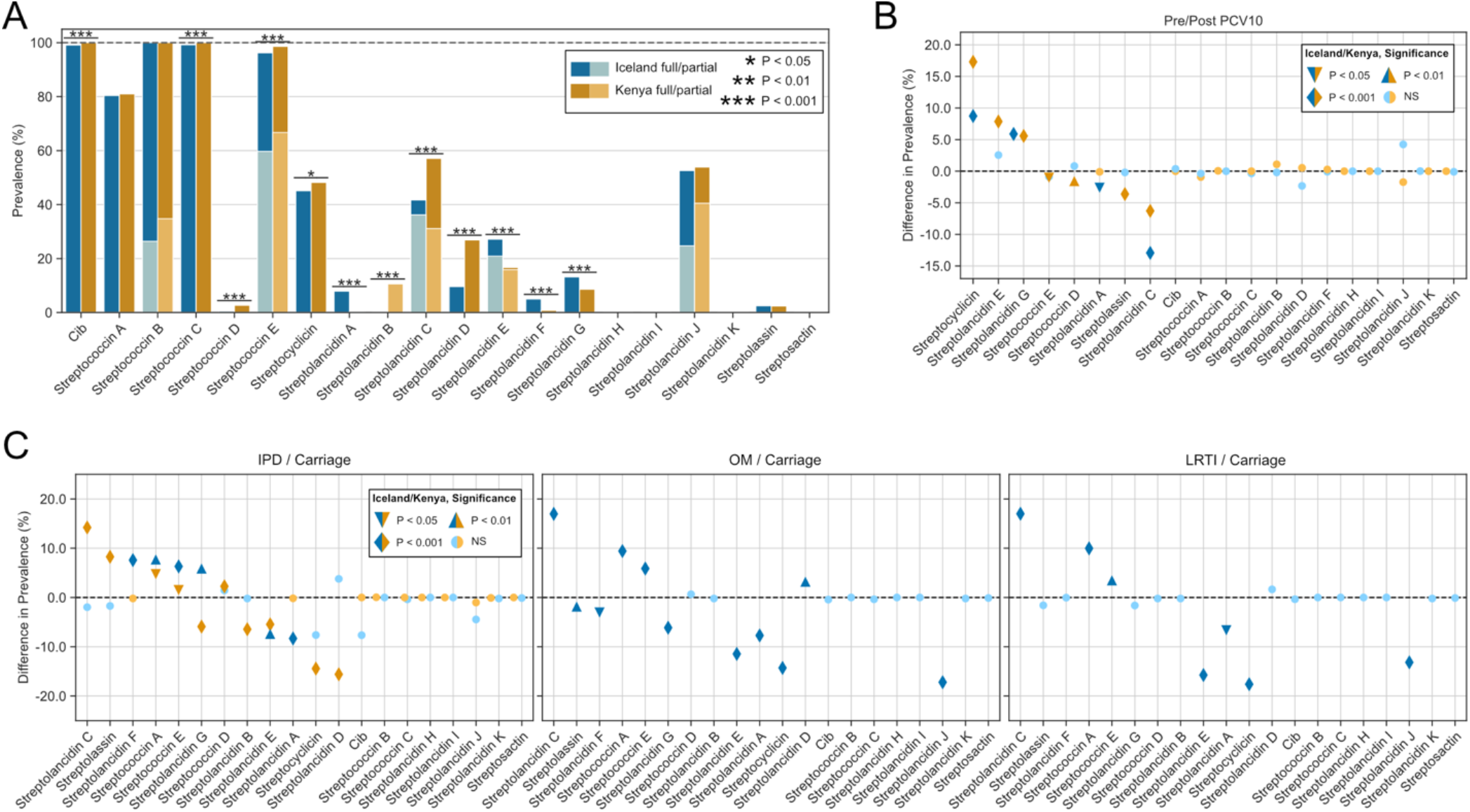
Prevalence of 19 different bacteriocin clusters in the Icelandic and Kenyan datasets. (A) Overall prevalence of each bacteriocin cluster in Iceland and Kenya. (B) Differences in prevalence of bacteriocin clusters in the post-PCV10 vs pre-PCV10 time periods. (C) Differences in the prevalence of bacteriocin clusters among invasive pneumococci (IPD) vs carriage pneumococci in Iceland and Kenya (left panel), pneumococci causing otitis media (OM) vs carriage pneumococci in Iceland (middle panel), and pneumococci causing lower respiratory tract infection (LRTI) vs carriage pneumococci in Iceland (right panel). Icelandic data are displayed with blue bars and symbols, and Kenyan data are displayed with tan bars and symbols. Significant differences were assessed using a Chi-square test. See Methods for definition of full and partial clusters.

### Bacteriocin prevalence was altered in the post-PCV10 period

Statistically significant differences in the prevalence of eight bacteriocin clusters were observed among pneumococci in the post-PCV10 period (Figure 2B). Among Icelandic pneumococci the prevalence of streptocyclicin and streptolancidin G increased, and streptolancidins A and C decreased. Among Kenyan pneumococci, streptocyclicin and streptolancidins E and G increased in prevalence, and streptococcins E and D, streptolancidin C and streptolassin decreased in prevalence.

### Bacteriocin cluster prevalence differed among pneumococci from carriage and disease

A comparison of carriage and invasive Kenyan pneumococci revealed significant differences in the prevalence of 10 bacteriocin clusters, five of which were more common among invasive pneumococci, and five were more common among carriage pneumococci (Figure 2C, left panel). A range of serotypes were represented among invasive and carriage pneumococci (Supplementary Table 7). Similarly, significant differences in the prevalence of bacteriocin clusters were observed among Icelandic pneumococci from disease and carriage: IPD vs carriage, six bacteriocins, OM vs carriage, 11 bacteriocins; and LRTI vs carriage, seven bacteriocins (Figure 2C, all three panels). Overall, streptococcin A and E clusters were significantly more common among Icelandic pneumococci recovered from all three disease processes relative to carriage pneumococci, and a range of serotypes were represented (Figure 2C; Supplementary Table 7).

### Bacteriocin cluster prevalence and post-vaccine population restructuring

The observed differences in bacteriocin prevalence were investigated relative to changes in the frequency of CCs pre- and post-PCV10 implementation. For example, streptolancidin C clusters were significantly associated with pre-PCV10 pneumococci in both datasets (Figure 2B), specifically with pneumococci causing IPD in the Kenyan dataset, and OM and LRTI in the Icelandic dataset (Figure 2C). Streptolancidin C clusters were mostly associated with pneumococci of CCs associated with PCV10 serotypes (Figure 3A).

**Figure 3:**
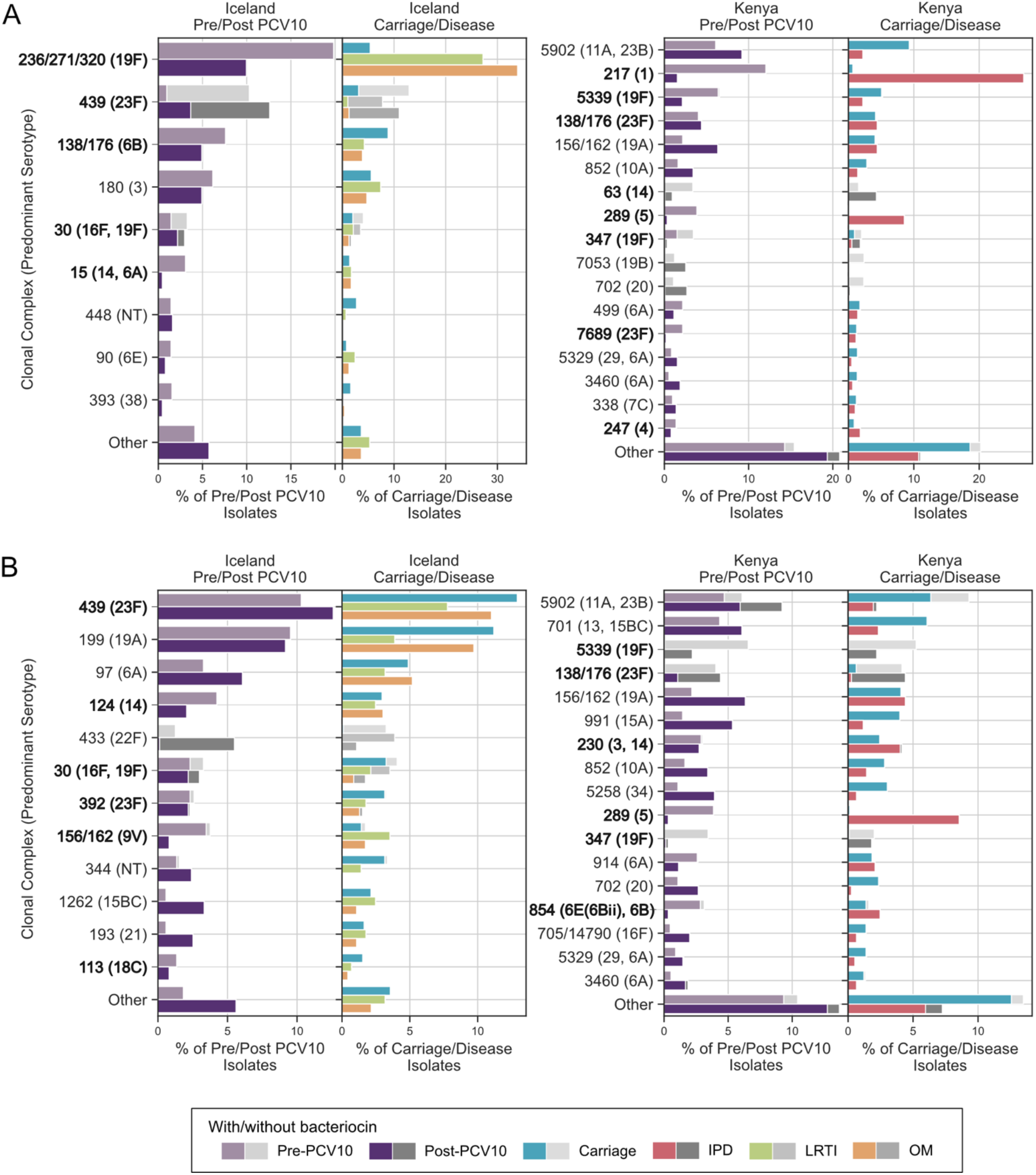
Prevalence of streptolancidin C (panel A) and streptocyclicin (panel B) bacteriocin clusters among pneumococci in two groups, carriage vs disease and pre- vs post-PCV10, stratified by clonal complex (CC). Each plot shows all CCs in which the bacteriocin cluster was detected. Any CC representing <1% of the overall dataset was in the ‘Other’ category. Each bar represents the overall percentage of pneumococci within that CC, the coloured section of the bar represents those pneumococci with the bacteriocin cluster, and the grey section represents those without the bacteriocin cluster. For Icelandic disease-causing pneumococci, only the disease in which the bacteriocin cluster was significantly altered relative to carriage are shown. Note: The dominant serotype(s) of each CC is shown in brackets. PCV10 serotypes are in bold font. IPD, invasive pneumococcal disease; LRTI, lower respiratory tract infection; OM, otitis media.

In contrast, streptocyclicin clusters were significantly more prevalent post-PCV10 (Figure 2B), among carriage rather than disease pneumococci (Figure 2C), and were common among both vaccine and nonvaccine serotype pneumococci (Figure 3B). All of the bacteriocin clusters with significantly different pre- or post-PCV10 frequencies (Figure 2B) were inspected, and changes in the prevalence of CCs pre-/post-PCV10 introduction were generally found to explain the differences in bacteriocin prevalence (Supplementary Tables 8 and 9).

### Bacteriocin repertoires shared between the two datasets and within lineages

Finally, the combination of different bacteriocin clusters in each genome, or ‘repertoire’, was investigated. Overall, each pneumococcal genome harboured between 4 and 11 different bacteriocin clusters, and in both datasets the mode was seven (Figure 4A). 89% and 81% of genomes had between 6 and 8 bacteriocin clusters in the Icelandic and Kenyan datasets, respectively. Overall, 134 different bacteriocin repertoires were observed, although more variation was observed in the Kenyan dataset (n=103 repertoires) than the Icelandic dataset (n=74 repertoires). In total, 43 repertoires were common to both datasets and the most frequently observed one included cib, streptococcins A, B, C, E, and streptolancidin C (Figure 4B).

**Figure 4:**
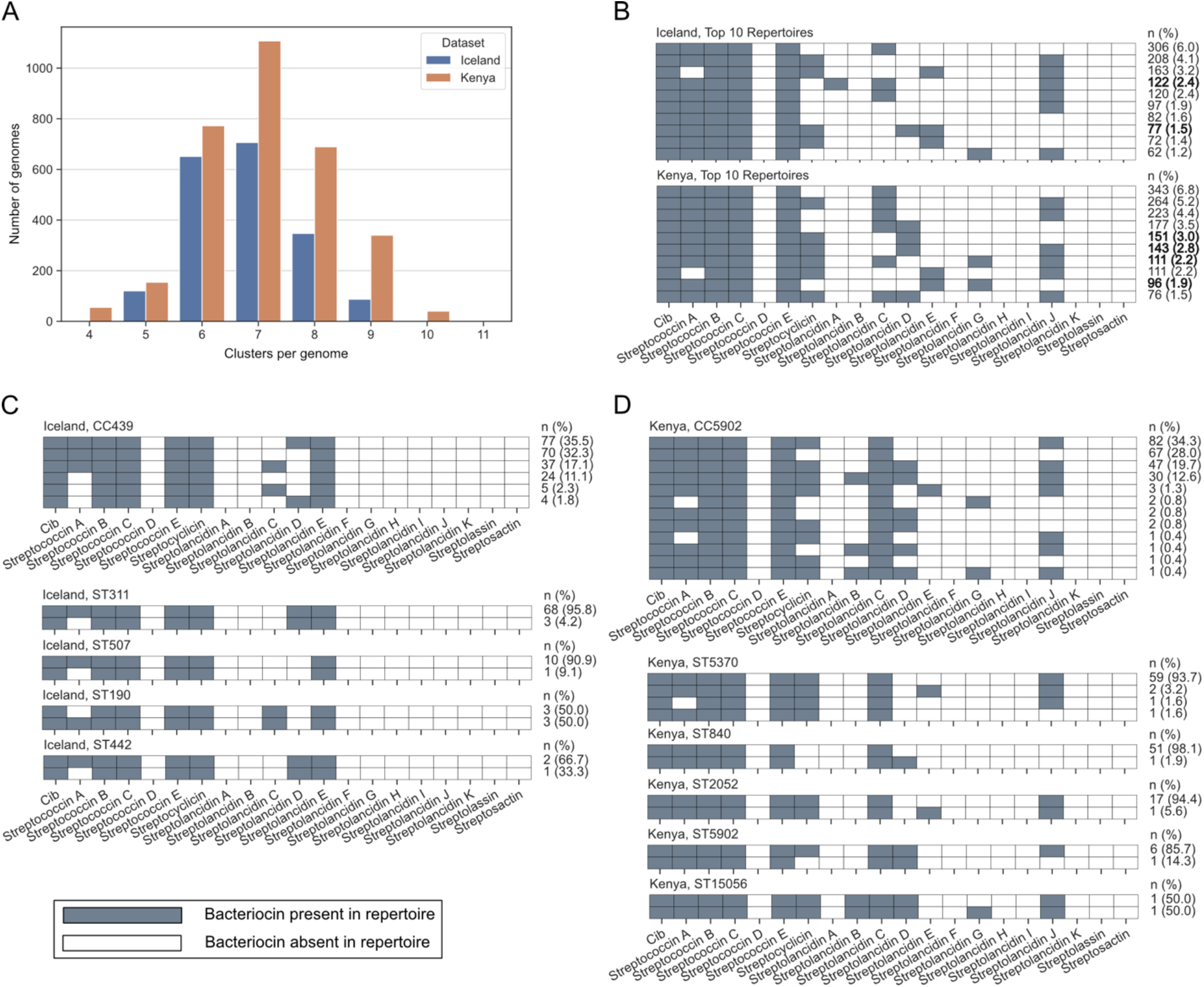
Bacteriocin repertoires observed among Icelandic and Kenyan pneumococci. (A) Number of bacteriocin clusters detected per genome. (B) The composition of the 10 most frequently observed bacteriocin repertoires in Icelandic and Kenyan pneumococci. Bold text indicates that the repertoire was found only in that country. (C, D) Bacteriocin repertoires observed in genomes from CC439 in the Icelandic dataset (panel C), and CC5902 in the Kenyan dataset (panel D), including a more detailed breakdown of the CC by sequence type (ST) below the CC summary. STs with no differences in repertoire are not shown. Note: the frequency of each repertoire within the Icelandic or Kenyan dataset, respectively, is given at the far right of each diagram.

The bacteriocin repertoire was generally consistent among all pneumococci within the same CC; although there were examples of CCs (Iceland, n=29; Kenya, n=61) in which more than one bacteriocin repertoire was observed and some of the STs that comprised the CC also had variable repertoires (Supplementary Tables 10 and 11). For example, CC439 in Iceland (Figure 4C) and CC5902 (Figure 4D) in Kenya demonstrated minor differences in the presence or absence of bacteriocin clusters, and these differences were maintained even within some STs.

## Discussion

Bacteriocins are complicated in terms of their high prevalence and distribution among pneumococci, but there are also clear patterns emerging from this and previously published work that allow for more focused study of individual bacteriocin clusters, repertoires of bacteriocins, and the pneumococcal lineages that harbour them. In this study we observed significant differences in bacteriocin cluster prevalence between Icelandic and Kenyan pneumococci, carriage and disease pneumococci, and pneumococci recovered pre- and post-PCV10 introduction. These observations could largely be explained by different underlying pneumococcal population structures in each country, as bacteriocin clusters tended to be associated with specific CCs. Nevertheless, a bacteriocin repertoire was not necessarily fixed within a CC and there were many examples where genetically related pneumococci had gained or lost bacteriocin clusters. This would be consistent with horizontal gene transfer of whole bacteriocin clusters among pneumococci, either by homologous recombination, or within integrative conjugative elements (ICEs)^33-36^.

These patterns are important because they provide the framework on which to propose hypotheses and experiments that allow for further study of these bacteriocins. It is also essential to understand any individual bacteriocin clusters in the context of all the other bacteriocin clusters present within the same genome, rather than in isolation, since every pneumococcal genome possesses many different bacteriocin clusters. Notably, our earlier study showed that multiple bacteriocin genes from different bacteriocin clusters were expressed simultaneously, but how the expression and regulation of these clusters is governed is still unknown^14^.

An additional factor in this complexity is that some of these bacteriocin clusters are also detectable in genomes of non-pneumococcal streptococci such as *Streptococcus mitis, Streptococcus oralis* and *Streptococcus pseuodopneumoniae*^14^. Horizontal genetic exchange between pneumococci and non-pneumococcal streptococci has been documented since the 1990s and is a major driver in the evolution of these organisms^37-40^. Understanding bacteriocin-mediated competition between different *Streptococcus* species (and other nasopharyngeal colonisers) will also be required to fully understand microbial interactions in the nasopharynx.

Moreover, if pneumococcal bacteriocin clusters are horizontally exchanged among pneumococci and/or other streptococci, one consequence is that genetic lineages could adapt to altered competition dynamics in remodelled post-PCV populations by acquiring a bacteriocin repertoire that improves competitiveness, and this may (or may not) lead to increased pneumococcal disease. In this study it was not possible to consider the dynamics of bacteriocins among co-colonising pneumococcal strains, nor could we investigate contributions from other commensal microbes that share the same ecological niche and presumably compete for the same resources. Experimental studies of microbial competition would likely be helpful in this context.

Overall, this study focused on bacteriocins in two large, geographically distinct, well characterised pneumococcal populations, and revealed the prevalence and diversity of bacteriocins harboured by carriage and disease-causing pneumococci with greater clarity. This work also demonstrated that PCV-mediated perturbations in the pneumococcal population structure led to changes in the distribution of bacteriocin clusters, which may have consequences for pneumococcal disease. Further work will be required to establish the role of pneumococcal bacteriocins in nasopharyngeal competition and also the range of species targeted by these bacteriocins.

## Data availability

Analysis code is available at GitHub: https://github.com/brueggemann-lab/bacteriocins_IceKen_2022. Assembled genomes will be publicly available in PubMLST upon publication (https://pubmlst.org/organisms/streptococcus-pneumoniae; Supplementary Tables 1 and 2).

## Acknowledgments

The authors are grateful to Dr Keith Jolley for PubMLST database support.

## Author contributions

MEBB, MJJvR and ABB led the study, managed and curated the data, and performed the data analyses. AK, BM, DA, SJQ, GH, HE, AH, KGK and JAGS performed the studies in Iceland and Kenya that led to the large collections of isolates characterised in this study. AA, CF and AJvT prepared DNA extracts for sequencing. SDB managed the genome sequencing. MEBB and ABB wrote the manuscript and all authors contributed to it before submission.

## References

1. GBD 2016 Lower Respiratory Infections Collaborators. Estimates of the global, regional, and national morbidity, mortality, and aetiologies of lower respiratory infections in 195 countries, 1990-2016: a systematic analysis for the Global Burden of Disease Study 2016. Lancet Infect Dis. 2018 Nov;18(11):1191–1210. doi: 10.1016/S1473-3099(18)30310-4. Epub 2018 Sep 19. PMID: 30243584; PMCID: PMC6202443.

2. Tuomanen EI, Mitchell TJ, Morrison DA, Spratt BG. The Pneumococcus. ASM Press; 2004. doi:10.1128/9781555816537.

3. Sleeman KL, Griffiths D, Shackley F, Diggle L, Gupta S, Maiden MC, Moxon ER, Crook DW, Peto TE. Capsular serotype-specific attack rates and duration of carriage of Streptococcus pneumoniae in a population of children. J Infect Dis. 2006 Sep 1;194(5):682–8. doi: 10.1086/505710. Epub 2006 Jul 25. PMID: 16897668.

4. Shak JR, Vidal JE, Klugman KP. Influence of bacterial interactions on pneumococcal colonization of the nasopharynx. Trends Microbiol. 2013 Mar;21(3):129–35. doi: 10.1016/j.tim.2012.11.005. Epub 2012 Dec 25. PMID: 23273566; PMCID: PMC3729046.

5. Auranen K, Mehtälä J, Tanskanen A, S Kaltoft M. Between-strain competition in acquisition and clearance of pneumococcal carriage--epidemiologic evidence from a longitudinal study of day-care children. Am J Epidemiol. 2010 Jan 15;171(2):169–76. doi: 10.1093/aje/kwp351. Epub 2009 Dec 6. PMID: 19969530; PMCID: PMC2800239.

6. Hammitt LL, Etyang AO, Morpeth SC, Ojal J, Mutuku A, Mturi N, Moisi JC, Adetifa IM, Karani A, Akech DO, Otiende M, Bwanaali T, Wafula J, Mataza C, Mumbo E, Tabu C, Knoll MD, Bauni E, Marsh K, Williams TN, Kamau T, Sharif SK, Levine OS, Scott JAG. Effect of ten-valent pneumococcal conjugate vaccine on invasive pneumococcal disease and nasopharyngeal carriage in Kenya: a longitudinal surveillance study. Lancet. 2019 May 25;393(10186):2146–2154. doi: 10.1016/S0140-6736(18)33005-8. Epub 2019 Apr 15. PMID: 31000194; PMCID: PMC6548991.

7. Quirk SJ, Haraldsson G, Erlendsdóttir H, Hjálmarsdóttir MÁ, van Tonder AJ, Hrafnkelsson B, Sigurdsson S, Bentley SD, Haraldsson Á, Brueggemann AB, Kristinsson KG. Effect of vaccination on pneumococci isolated from the nasopharynx of healthy children and the middle ear of children with otitis media in Iceland. J Clin Microbiol. 2018 Nov 27;56(12):e01046–18. doi: 10.1128/JCM.01046-18. PMID: 30257906; PMCID: PMC6258863.

8. Quirk SJ, Haraldsson G, Hjálmarsdóttir MÁ, van Tonder AJ, Hrafnkelsson B, Bentley SD, Haraldsson Á, Erlendsdóttir H, Brueggemann AB, Kristinsson KG. Vaccination of Icelandic children with the 10-valent pneumococcal vaccine leads to a significant herd effect among adults in Iceland. J Clin Microbiol. 2019 Mar 28;57(4):e01766–18. doi: 10.1128/JCM.01766-18. PMID: 30651396; PMCID: PMC6440763.

9. Dawid S, Roche AM, Weiser JN. The blp bacteriocins of Streptococcus pneumoniae mediate intraspecies competition both in vitro and in vivo. Infect Immun. 2007 Jan;75(1):443–51. doi: 10.1128/IAI.01775-05. Epub 2006 Oct 30. PMID: 17074857; PMCID: PMC1828380.

10. Perez RH, Zendo T, Sonomoto K. Novel bacteriocins from lactic acid bacteria (LAB): various structures and applications. Microb Cell Fact. 2014 Aug 29;13 Suppl 1(Suppl 1):S3. doi: 10.1186/1475-2859-13-S1-S3. Epub 2014 Aug 29. PMID: 25186038; PMCID: PMC4155820.

11. Arnison PG, Bibb MJ, Bierbaum G, Bowers AA, Bugni TS, Bulaj G, Camarero JA, Campopiano DJ, Challis GL, Clardy J, Cotter PD, Craik DJ, Dawson M, Dittmann E, Donadio S, Dorrestein PC, Entian KD, Fischbach MA, Garavelli JS, Göransson U, Gruber CW, Haft DH, Hemscheidt TK, Hertweck C, Hill C, Horswill AR, Jaspars M, Kelly WL, Klinman JP, Kuipers OP, Link AJ, Liu W, Marahiel MA, Mitchell DA, Moll GN, Moore BS, Müller R, Nair SK, Nes IF, Norris GE, Olivera BM, Onaka H, Patchett ML, Piel J, Reaney MJ, Rebuffat S, Ross RP, Sahl HG, Schmidt EW, Selsted ME, Severinov K, Shen B, Sivonen K, Smith L, Stein T, Süssmuth RD, Tagg JR, Tang GL, Truman AW, Vederas JC, Walsh CT, Walton JD, Wenzel SC, Willey JM, van der Donk WA. Ribosomally synthesized and post-translationally modified peptide natural products: overview and recommendations for a universal nomenclature. Nat Prod Rep. 2013 Jan;30(1):108–60. doi: 10.1039/c2np20085f. PMID: 23165928; PMCID: PMC3954855.

12. Bogaardt C, van Tonder AJ, Brueggemann AB. Genomic analyses of pneumococci reveal a wide diversity of bacteriocins - including pneumocyclicin, a novel circular bacteriocin. BMC Genomics. 2015 Jul 28;16(1):554. doi: 10.1186/s12864-015-1729-4. PMID: 26215050; PMCID: PMC4517551.

13. Wholey WY, Abu-Khdeir M, Yu EA, Siddiqui S, Esimai O, Dawid S. Characterization of the competitive pneumocin peptides of Streptococcus pneumoniae. Front Cell Infect Microbiol. 2019 Mar 12;9:55. doi: 10.3389/fcimb.2019.00055. PMID: 30915281; PMCID: PMC6422914.

14. Rezaei Javan R, van Tonder AJ, King JP, Harrold CL, Brueggemann AB. Genome sequencing reveals a large and diverse repertoire of antimicrobial peptides. Front Microbiol. 2018 Aug 27;9:2012. doi: 10.3389/fmicb.2018.02012. PMID: 30210481; PMCID: PMC6120550.

15. Begley M, Cotter PD, Hill C, Ross RP. Identification of a novel two-peptide lantibiotic, lichenicidin, following rational genome mining for LanM proteins. Appl Environ Microbiol. 2009 Sep;75(17):5451–60. doi: 10.1128/AEM.00730-09. Epub 2009 Jun 26. PMID: 19561184; PMCID: PMC2737927.

16. Abdullahi O, Nyiro J, Lewa P, Slack M, Scott JA. The descriptive epidemiology of Streptococcus pneumoniae and Haemophilus influenzae nasopharyngeal carriage in children and adults in Kilifi district, Kenya. Pediatr Infect Dis J. 2008 Jan;27(1):59–64. doi: 10.1097/INF.0b013e31814da70c. PMID: 18162940; PMCID: PMC2382474.

17. Abdullahi O, Karani A, Tigoi CC, Mugo D, Kungu S, Wanjiru E, Jomo J, Musyimi R, Lipsitch M, Scott JA. The prevalence and risk factors for pneumococcal colonization of the nasopharynx among children in Kilifi District, Kenya. PLoS One. 2012;7(2):e30787. doi: 10.1371/journal.pone.0030787. Epub 2012 Feb 20. PMID: 22363489; PMCID: PMC3282706.

18. Abdullahi O, Karani A, Tigoi CC, Mugo D, Kungu S, Wanjiru E, Jomo J, Musyimi R, Lipsitch M, Scott JA. Rates of acquisition and clearance of pneumococcal serotypes in the nasopharynges of children in Kilifi District, Kenya. J Infect Dis. 2012 Oct 1;206(7):1020–9. doi: 10.1093/infdis/jis447. Epub 2012 Jul 24. PMID: 22829650; PMCID: PMC3433858.

19. Hammitt LL, Akech DO, Morpeth SC, Karani A, Kihuha N, Nyongesa S, Bwanaali T, Mumbo E, Kamau T, Sharif SK, Scott JA. Population effect of 10-valent pneumococcal conjugate vaccine on nasopharyngeal carriage of Streptococcus pneumoniae and non-typeable Haemophilus influenzae in Kilifi, Kenya: findings from cross-sectional carriage studies. Lancet Glob Health. 2014 Jul;2(7):e397–405. doi: 10.1016/S2214-109X(14)70224-4. Epub 2014 May 28. PMID: 25103393; PMCID: PMC5628631.

20. van Tonder AJ, Bray JE, Roalfe L, White R, Zancolli M, Quirk SJ, Haraldsson G, Jolley KA, Maiden MC, Bentley SD, Haraldsson Á, Erlendsdóttir H, Kristinsson KG, Goldblatt D, Brueggemann AB. Genomics reveals the worldwide distribution of multidrug-resistant serotype 6E pneumococci. J Clin Microbiol. 2015 Jul;53(7):2271–85. doi: 10.1128/JCM.00744-15. Epub 2015 May 13.

21. van Tonder AJ, Bray JE, Quirk SJ, Haraldsson G, Jolley KA, Maiden MCJ, Hoffmann S, Bentley SD, Haraldsson Á, Erlendsdóttir H, Kristinsson KG, Brueggemann AB. Putatively novel serotypes and the potential for reduced vaccine effectiveness: capsular locus diversity revealed among 5405 pneumococcal genomes. Microb Genom. 2016 Oct 1;2(10):000090. doi: 10.1099/mgen.0.000090. PMID: 28133541; PMCID: PMC5266551.

22. Zerbino DR. Using the Velvet de novo assembler for short-read sequencing technologies. Curr Protoc Bioinformatics. 2010 Sep;Chapter 11:Unit 11.5. doi: 10.1002/0471250953.bi1105s31. PMID: 20836074; PMCID: PMC2952100.

23. Boetzer M, Henkel CV, Jansen HJ, Butler D, Pirovano W. Scaffolding pre-assembled contigs using SSPACE. Bioinformatics. 2011 Feb 15;27(4):578–9. doi: 10.1093/bioinformatics/btq683. Epub 2010 Dec 12. PMID: 21149342.

24. Boetzer M, Pirovano W. Toward almost closed genomes with GapFiller. Genome Biol. 2012 Jun 25;13(6):R56. doi: 10.1186/gb-2012-13-6-r56. PMID: 22731987; PMCID: PMC3446322.

25. Jolley KA, Maiden MC. BIGSdb: Scalable analysis of bacterial genome variation at the population level. BMC Bioinformatics. 2010 Dec 10;11:595. doi: 10.1186/1471-2105-11-595. PMID: 21143983; PMCID: PMC3004885.

26. Jolley KA, Bliss CM, Bennett JS, Bratcher HB, Brehony C, Colles FM, Wimalarathna H, Harrison OB, Sheppard SK, Cody AJ, Maiden MCJ. Ribosomal multilocus sequence typing: universal characterization of bacteria from domain to strain. Microbiology (Reading). 2012 Apr;158(Pt 4):1005–1015. doi: 10.1099/mic.0.055459-0. Epub 2012 Jan 27. PMID: 22282518; PMCID: PMC3492749.

27. Francisco AP, Vaz C, Monteiro PT, Melo-Cristino J, Ramirez M, Carriço JA. PHYLOViZ: phylogenetic inference and data visualization for sequence based typing methods. BMC Bioinformatics. 2012 May 8;13:87. doi: 10.1186/1471-2105-13-87. PMID: 22568821; PMCID: PMC3403920.

28. Epping L, van Tonder AJ, Gladstone RA, The Global Pneumococcal Sequencing Consortium, Bentley SD, Page AJ, Keane JA. SeroBA: rapid high-throughput serotyping of Streptococcus pneumoniae from whole genome sequence data. Microb Genom. 2018 Jul;4(7):e000186. doi: 10.1099/mgen.0.000186. Epub 2018 Jun 15.

29. Guiral S, Mitchell TJ, Martin B, Claverys JP. Competence-programmed predation of noncompetent cells in the human pathogen Streptococcus pneumoniae: genetic requirements. Proc Natl Acad Sci U S A. 2005 Jun 14;102(24):8710–5. doi: 10.1073/pnas.0500879102. Epub 2005 May 31. PMID: 15928084; PMCID: PMC1150823.

30. Hoover SE, Perez AJ, Tsui HC, Sinha D, Smiley DL, DiMarchi RD, Winkler ME, Lazazzera BA. A new quorum-sensing system (TprA/PhrA) for Streptococcus pneumoniae D39 that regulates a lantibiotic biosynthesis gene cluster. Mol Microbiol. 2015 Jul;97(2):229–43. doi: 10.1111/mmi.13029. Epub 2015 May 26. PMID: 25869931; PMCID: PMC4676566.

31. Maricic N, Anderson ES, Opipari AE, Yu EA, Dawid S. Characterization of a multipeptide lantibiotic locus in Streptococcus pneumoniae. mBio. 2016 Jan 26;7(1):e01656–15. doi: 10.1128/mBio.01656-15. PMID: 26814178; PMCID: PMC4742701.

32. Walker GV, Heng NCK, Carne A, Tagg JR, Wescombe PA. Salivaricin E and abundant dextranase activity may contribute to the anti-cariogenic potential of the probiotic candidate Streptococcus salivarius JH. Microbiology (Reading). 2016 Mar;162(3):476–486. doi: 10.1099/mic.0.000237. Epub 2016 Jan 7. PMID: 26744310.

33. Croucher NJ, Walker D, Romero P, Lennard N, Paterson GK, Bason NC, Mitchell AM, Quail MA, Andrew PW, Parkhill J, Bentley SD, Mitchell TJ. Role of conjugative elements in the evolution of the multidrug-resistant pandemic clone Streptococcus pneumoniae Spain23F ST81. J Bacteriol. 2009 Mar;191(5):1480–9. doi: 10.1128/JB.01343-08. Epub 2008 Dec 29. PMID: 19114491; PMCID: PMC2648205.

34. Wyres KL, van Tonder A, Lambertsen LM, Hakenbeck R, Parkhill J, Bentley SD, Brueggemann AB. Evidence of antimicrobial resistance-conferring genetic elements among pneumococci isolated prior to 1974. BMC Genomics. 2013 Jul 24;14:500. doi: 10.1186/1471-2164-14-500. PMID: 23879707; PMCID: PMC3726389.

35. Ambroset C, Coluzzi C, Guédon G, Devignes MD, Loux V, Lacroix T, Payot S, Leblond-Bourget N. New insights into the classification and integration specificity of Streptococcus integrative conjugative elements through extensive genome exploration. Front Microbiol. 2016 Jan 6;6:1483. doi: 10.3389/fmicb.2015.01483. PMID: 26779141; PMCID: PMC4701971.

36. Sun Y, Veseli IA, Vaillancourt K, Frenette M, Grenier D, Pombert JF. The bacteriocin from the prophylactic candidate Streptococcus suis 90-1330 is widely distributed across S. suis isolates and appears encoded in an integrative and conjugative element. PLoS One. 2019 Apr 30;14(4):e0216002. doi: 10.1371/journal.pone.0216002. PMID: 31039174; PMCID: PMC6490898.

37. Dowson CG, Hutchison A, Woodford N, Johnson AP, George RC, Spratt BG. Penicillin-resistant viridans streptococci have obtained altered penicillin-binding protein genes from penicillin-resistant strains of Streptococcus pneumoniae. Proc Natl Acad Sci U S A. 1990 Aug;87(15):5858–62. doi: 10.1073/pnas.87.15.5858. PMID: 2377622; PMCID: PMC54428.

38. Laible G, Spratt BG, Hakenbeck R. Interspecies recombinational events during the evolution of altered PBP 2x genes in penicillin-resistant clinical isolates of Streptococcus pneumoniae. Mol Microbiol. 1991 Aug;5(8):1993–2002. doi: 10.1111/j.1365-2958.1991.tb00821.x. PMID: 1766375.

39. Enright MC, Spratt BG. Extensive variation in the ddl gene of penicillin-resistant Streptococcus pneumoniae results from a hitchhiking effect driven by the penicillin-binding protein 2b gene. Mol Biol Evol. 1999 Dec;16(12):1687–95. doi: 10.1093/oxfordjournals.molbev.a026082. PMID: 10605111.

40. Salvadori G, Junges R, Morrison DA, Petersen FC. Competence in Streptococcus pneumoniae and close commensal relatives: mechanisms and implications. Front Cell Infect Microbiol. 2019 Apr 3;9:94. doi: 10.3389/fcimb.2019.00094. PMID: 31001492; PMCID: PMC6456647.

