## Supplementary_Tables_Figure for "Nasopharyngeal competition dynamics are likely to be altered following vaccine introduction: bacteriocin prevalence and diversity among Icelandic and Kenyan pneumococci"

### Supplementary data

Genome and isolate metadata released at time of publication:

**Supplementary Table 1**: PubMLST identification numbers of genomes included in the final Kenyan genomic dataset

**Supplementary Table 2**: PubMLST identification numbers of genomes included in the final Icelandic genomic dataset, including four that were excluded from original dataset (id numbers 1586, 1594, 1644, 2540)

**Supplementary Table 3: Genes associated with bacteriocin clusters in the Icelandic and Kenyan datasets.**

| Bacteriocin | Bacteriocin type | Gene | Typical length (bp) [a] | Predicted functionality |
|---|---|---|---|---|
| Cib[b,c] | Competence-induced | *cibA* | 186 | Bacteriocin precursor |
| | | *cibB* | 153 | Bacteriocin precursor |
| | | *cibC* | 198 | Immunity |
| Streptococcin A[b] | Lactococcin 972-like | *scaA* | 285 | Bacteriocin precursor |
| | | *scaB* | 2109 | Immunity |
| | | *scaC* | 642 | Immunity |
| Streptococcin B[b] | Lactococcin 972-like | *scbA* | 297 | Bacteriocin precursor |
| | | *scbB* | 2031 | Immunity |
| | | *scbC* | 642 | Immunity |
| Streptococcin C[b] | Lactococcin 972-like | *sccA* | 348 | Bacteriocin precursor |
| | | *sccB* | 2022 | Immunity |
| | | *sccC* | 642 | Immunity |
| Streptococcin D[b] | Lactococcin 972-like | *scdA* | 297 | Bacteriocin precursor |
| | | *scdB* | 2010 | Immunity |
| | | *scdC* | 633 | Immunity |
| Streptococcin E[b] | Lactococcin 972-like | *sceA* | 297 | Bacteriocin precursor |
| | | *sceB* | 2016 | Immunity |
| | | *sceC* | 633 | Immunity |
| Streptocyclicin[b,d] | Head-to-tail cyclised peptides | *scyA* | 297 | Bacteriocin precursor |
| | | *scyB* | 1137 | Bacteriocin biosynthesis |
| | | *scyC* | 483 | Bacteriocin biosynthesis |
| | | *scyD* | 597 | Bacteriocin biosynthesis |
| | | *scyE* | 492 | Bacteriocin biosynthesis |
| Streptolancidin A[b,e,f] | Class II lanthipeptide | *slaA1* | 183 | Bacteriocin precursor |
| | | *slaA2* | 183 | Bacteriocin precursor |
| | | *slaA3* | 183 | Bacteriocin precursor |
| | | *slaA4* | 177 | Bacteriocin precursor |
| | | *slaA5* | 108 | Bacteriocin precursor |
| | | *slaF* | 738 | Immunity |
| | | *slaE* | 2016 | Immunity |
| | | *slaK* | 1575 | Histidine kinase/response regulator |
| | | *slaR* | 597 | Histidine kinase/response regulator |

| | | | | |
|---|---|---|---|---|
| | | *slaM* | 2955 | Bacteriocin biosynthesis |
| | | *slaT* | 2124 | Transporter |
| Streptolancidin B[b,g] | Class II lanthipeptide | *slbF* | 936 | Immunity |
| | | *slbG* | 741 | Immunity |
| | | *slbE* | 729 | Immunity |
| | | *slbA* | 216 | Bacteriocin precursor |
| | | *slbM* | 3252 | Bacteriocin biosynthesis |
| | | *slbT* | 2067 | Transporter |
| Streptolancidin C[b] | Class IV lanthipeptide | *slcA* | 105 | Bacteriocin precursor |
| | | *slcX* | 993 | Unknown |
| | | *slcL* | 1437 | Bacteriocin biosynthesis |
| | | *slcT* | 1239 | Transporter |
| Streptolancidin D[b] | Class I lanthipeptide | *sldA* | 105 | Bacteriocin precursor |
| | | *sldB* | 2793 | Bacteriocin biosynthesis |
| | | *sldC* | 1278 | Bacteriocin biosynthesis |
| | | *sldT* | 1257 | Transporter |
| Streptolancidin E[b,h] | Class II lanthipeptide | *sleM1* | 3036 | Bacteriocin biosynthesis |
| | | *sleA1* | 171 | Bacteriocin precursor |
| | | *sleA2* | 192 | Bacteriocin precursor |
| | | *sleM2* | 2016 | Bacteriocin biosynthesis |
| | | *sleM3* | 747 | Bacteriocin biosynthesis |
| | | *sleT* | 2142 | Transporter |
| | | *sleX1* | 171 | Unknown |
| | | *sleF* | 729 | Immunity |
| | | *sleG* | 738 | Immunity |
| | | *sleX2* | 711 | Unknown |
| Streptolancidin F[b] | Class IV lanthipeptide | *slfA* | 99 | Bacteriocin precursor |
| | | *slfL* | 2496 | Bacteriocin biosynthesis |
| Streptolancidin G[b,i] | Class II lanthipeptide | *slgA1* | 225 | Bacteriocin precursor |
| | | *slgA2* | 189 | Bacteriocin precursor |
| | | *slgM* | 2991 | Bacteriocin biosynthesis |
| | | *slgD* | 705 | Bacteriocin biosynthesis |
| | | *slgP1* | 930 | Bacteriocin biosynthesis |
| | | *slgT* | 2109 | Transporter |

| | | | | |
|---|---|---|---|---|
| | | *slgP2* | 1740 | Bacteriocin biosynthesis |
| Streptolancidin H[b] | Class I lanthipeptide | *slhP* | 1368 | Bacteriocin biosynthesis |
| | | *slhR* | 684 | Histidine kinase/response regulator |
| | | *slhK* | 1341 | Histidine kinase/response regulator |
| | | *slhF* | 687 | Immunity |
| | | *slhE* | 741 | Immunity |
| | | *slhG* | 678 | Immunity |
| | | *slhX1* | 645 | Unknown |
| | | *slhX2* | 285 | Unknown |
| | | *slhA* | 177 | Bacteriocin precursor |
| | | *slhB* | 2964 | Bacteriocin biosynthesis |
| | | *slhT* | 1776 | Transporter |
| | | *slhC* | 1272 | Bacteriocin biosynthesis |
| | | *slhI* | 663 | Immunity |
| Streptolancidin I[b] | Class I lanthipeptide | *sliP* | 1374 | Bacteriocin biosynthesis |
| | | *sliR* | 699 | Histidine kinase/response regulator |
| | | *sliK* | 1344 | Histidine kinase/response regulator |
| | | *sliF* | 702 | Immunity |
| | | *sliE* | 738 | Immunity |
| | | *sliG* | 687 | Immunity |
| | | *sliA* | 168 | Bacteriocin precursor |
| | | *sliB* | 2976 | Bacteriocin biosynthesis |
| | | *sliT* | 1809 | Transporter |
| | | *sliC* | 1278 | Bacteriocin biosynthesis |
| | | *sliI* | 717 | Immunity |
| Streptolancidin J[b] | Class IV lanthipeptide | *sljA1* | 138 | Bacteriocin precursor |
| | | *sljL* | 2610 | Bacteriocin biosynthesis |
| | | *sljP* | 1941 | Bacteriocin biosynthesis |
| | | *sljT1* | 1608 | Transporter |
| | | *sljT2* | 741 | Transporter |
| | | *sljT3* | 1320 | Transporter |

| | | | | |
|---|---|---|---|---|
| | | *sljA2* | 138 | Bacteriocin precursor |
| Streptolancidin K[b] | Class IV lanthipeptide | *slkA* | 99 | Bacteriocin precursor |
| | | *slkL* | 2517 | Bacteriocin biosynthesis |
| | | *slkT* | 1221 | Transporter |
| Streptolassin[b] | Lasso peptide | *slsA* | 129 | Bacteriocin precursor |
| | | *slsC* | 1725 | Bacteriocin biosynthesis |
| | | *slsB1* | 252 | Bacteriocin biosynthesis |
| | | *slsB2* | 2232 | Bacteriocin biosynthesis |
| | | *slsF* | 717 | Immunity |
| | | *slsE* | 792 | Immunity |
| | | *slsG* | 711 | Immunity |
| | | *slsR* | 774 | Histidine kinase/response regulator |
| | | *slsK* | 1098 | Histidine kinase/response regulator |
| Streptosactin[b] | Sactipeptide | *ssaA* | 177 | Bacteriocin precursor |
| | | *ssaCD* | 1338 | Bacteriocin biosynthesis |
| | | *ssaX1* | 153 | Unknown |
| | | *ssaX2* | 996 | Unknown |
| | | *ssaP* | 861 | Bacteriocin biosynthesis |
| | | *ssaX3* | 696 | Unknown |

a. Length of genes as published previously (see b)


b. Rezaei Javan R, van Tonder AJ, King JP, Harrold CL, Brueggemann AB. Genome sequencing reveals a large and diverse repertoire of antimicrobial peptides. *Front Microbiol*. 2018;9(AUG):1-15. doi:10.3389/fmicb.2018.02012

c. Guiral S, Mitchell TJ, Martin B, Claverys JP. Competence-programmed predation of noncompetent cells in the human pathogen *Streptococcus pneumoniae*: Genetic requirements. *Proc Natl Acad Sci U S A*. 2005;102(24):8710-8715. doi:10.1073/pnas.0500879102

d. Bogaardt C, van Tonder AJ, Brueggemann AB. Genomic analyses of pneumococci reveal a wide diversity of bacteriocins - including pneumocyclicin, a novel circular bacteriocin. *BMC Genomics*. 2015;16(1):554. doi:10.1186/s12864-015-1729-4

e. Maricic N, Anderson ES, Opipari AME, Yu EA, Dawid S. Characterization of a multipeptide lantibiotic locus in *Streptococcus pneumoniae*. *mBio*. 2016;7(1):1656-1671. doi:10.1128/mBio.01656-15

f. Walker G V., Heng NCK, Carne A, Tagg JR, Wescombe PA. Salivaricin E and abundant dextranase activity may contribute to the anti-cariogenic potential of the probiotic candidate *Streptococcus salivarius* JH. *Microbiol U K*. 2016;162(3):476-486. doi:10.1099/mic.0.000237

g. Kadam A, Eutsey RA, Rosch J, et al. Promiscuous signaling by a regulatory system unique to the pandemic PMEN1 pneumococcal lineage. *PLoS Pathog*. 2017;13(5). doi:10.1371/journal.ppat.1006339

h. Begley M, Cotter PD, Hill C, Ross RP. Identification of a novel two-peptide lantibiotic, lichenicidin, following rational genome mining for LanM proteins. *Appl Environ Microbiol*. 2009;75(17):5451-5460. doi:10.1128/AEM.00730-09

i. Hoover SE, Perez AJ, Tsui HCT, et al. A new quorum-sensing system (TprA/PhrA) for *Streptococcus pneumoniae* D39 that regulates a lantibiotic biosynthesis gene cluster. *Mol Microbiol*. 2015;97(2):229-243. doi:10.1111/mmi.1302

**Supplementary Table 4: Full, partial, and fragmented bacteriocin clusters identified among Icelandic and Kenyan pneumococci.**

| Bacteriocin | Country | Profile | Category | Frequency |
|---|---|---|---|---|
| Cib | Kenya | A-B-C | Full | 3159 |
| | Iceland | A-B-C | Full | 1895 |
| | | /-/-C | Fragment | 17 |
| Streptococcin A | Kenya | A-B-C | Full | 2562 |
| | Iceland | A-B-C | Full | 1537 |
| Streptococcin B | Kenya | A-B-C | Full | 2058 |
| | | /-B-C | Partial | 1100 |
| | | /-B-/ | Fragment | 1 |
| | Iceland | A-B-C | Full | 1408 |
| | | /-B-C | Partial | 504 |
| Streptococcin C | Kenya | A-B-C | Full | 3159 |
| | Iceland | A-B-C | Full | 1912 |
| Streptococcin D | Kenya | A-B-C | Full | 85 |
| | Iceland | A-B-C | Full | 9 |
| Streptococcin E | Kenya | A-B-C | Full | 1048 |
| | | /-B-C | Partial | 2107 |
| | Iceland | A-B-C | Full | 699 |
| | | /-B-C | Partial | 1142 |
| Streptocyclicin | Kenya | A-B-C-D-E | Full | 1524 |
| | | /-/-/-D-/ | Fragment | 1 |
| | Iceland | A-B-C-D-E | Full | 862 |
| Streptolancidin A | Kenya | A1-A2-A3-A4-A5-F-E-K-R-M-T | Full | 8 |
| | | /-/-/-/-/-/-/-/-/-/-T | Fragment | 2 |
| | Iceland | A1-A2-A3-A4-A5-F-E-K-R-M-T | Full | 152 |
| | | /-/-/-/-/-/-/-/-/-/-T | Fragment | 38 |
| Streptolancidin B | Kenya | F-G-E-A-M-T | Full | 2 |
| | | F-G-E-/-/-/ | Partial | 336 |
| | Iceland | F-G-E-/-/-/ | Partial | 2 |
| Streptolancidin C | Kenya | A-X-L-T | Full | 826 |
| | | A-X-/-/ | Partial | 983 |
| | | /-/-/-T | Fragment | 1 |
| | Iceland | A-X-L-T | Full | 106 |
| | | A-X-/-/ | Partial | 692 |
| Streptolancidin D | Kenya | A-B-C-T | Full | 853 |
| | Iceland | A-B-C-T | Full | 184 |
| Streptolancidin E | Kenya | M1-A1-A2-M2-M3-T-X1-F-G-X2 | Full | 29 |
| | | /-/-/-/-M3-T-X1-F-G-X2 | Partial | 493 |
| | | /-/-/-/-M3-T-/-F-G-X2 | Partial | 10 |

| | | | | |
|---|---|---|---|---|
| | Iceland | M1-A1-A2-M2-M3-T-X1-F-G-X2 | Full | 142 |
| | | /-/-/-/-M3-T-X1-F-G-X2 | Partial | 309 |
| | | /-/-/-/-M3-T-/-F-G-X2 | Partial | 91 |
| Streptolancidin F | Kenya | A-L | Full | 25 |
| | Iceland | A-L | Full | 95 |
| Streptolancidin G | Kenya | A1-A2-M-D-P1-T-P2 | Full | 272 |
| | Iceland | A1-A2-M-D-P1-T-P2 | Full | 252 |
| Streptolancidin J | Kenya | A1-L-P-T1-T2-T3-A2 | Full | 426 |
| | | A1-L-P-T1-T2-T3-/ | Partial | 1 |
| | | A1-L-/-T1-T2-T3-A2 | Partial | 931 |
| | | A1-L-/-T1-T2-T3-/ | Partial | 340 |
| | | A1-/-/-/-/-/-/ | Fragment | 2 |
| | | /-L-P-T1-T2-T3-A2 | Partial | 1 |
| | | /-L-/-T1-T2-T3-A2 | Partial | 6 |
| | | /-L-/-T1-T2-T3-/ | Partial | 1 |
| | | /-/-/-/-/-T3-/ | Fragment | 1 |
| | Iceland | A1-L-P-T1-T2-T3-A2 | Full | 535 |
| | | A1-L-P-T1-T2-T3-/ | Partial | 40 |
| | | A1-L-/-T1-T2-T3-A2 | Partial | 383 |
| | | A1-L-/-T1-T2-T3-/ | Partial | 30 |
| | | A1-L-/-/-T2-T3-A2 | Partial | 1 |
| | | /-L-P-T1-T2-T3-A2 | Partial | 12 |
| | | /-L-/-T1-T2-T3-A2 | Partial | 6 |
| Streptolancidin K | Kenya | A-L-T | Full | 2 |
| | | /-/-T | Fragment | 3 |
| | Iceland | A-L-T | Full | 2 |
| | | /-/-T | Fragment | 5 |
| Streptolassin | Kenya | A-C-B1-B2-F-E-G-R-K | Full | 77 |
| | Iceland | A-C-B1-B2-F-E-G-R-K | Full | 48 |
| Streptosactin | Iceland | A-CD-X1-X2-P-X3 | Full | 1 |

Note: Rows shaded in grey indicate fragmented clusters, which were excluded from further analysis.

**Supplementary Table 5: Full and partial bacteriocin clusters identified among Icelandic and Kenyan pneumococci.**

| **Iceland** | | | |
|---|---|---|---|
| **Bacteriocin cluster** | **Category** | **Frequency** | **% of total bacteriocin clusters** |
| Cib | Contiguous | 1895 | 100 |
| Streptococcin A | Contiguous | 1537 | 100 |
| Streptococcin B | Contiguous | 1912 | 100 |
| Streptococcin C | Contiguous | 1811 | 94.72 |
| | EOC | 82 | 4.29 |
| | Non-contiguous (multiple contigs, not EOC-adjacent) | 16 | 0.84 |
| | Contiguous with Ns | 3 | 0.16 |
| Streptococcin D | Contiguous | 9 | 100 |
| Streptococcin E | Contiguous | 1840 | 99.95 |
| | Non-contiguous (multiple contigs, not EOC-adjacent) | 1 | 0.05 |
| Streptocyclicin | Contiguous | 860 | 99.77 |
| | EOC | 2 | 0.23 |
| Streptolancidin A | Contiguous | 152 | 100 |
| Streptolancidin B | Contiguous | 2 | 100 |
| Streptolancidin C | Contiguous | 798 | 100 |
| Streptolancidin D | Contiguous | 183 | 99.46 |
| | EOC | 1 | 0.54 |
| Streptolancidin E | Contiguous | 461 | 85.06 |
| | EOC | 47 | 8.67 |
| | Non-contiguous (multiple contigs, not EOC-adjacent) | 15 | 2.77 |
| | Contiguous with Ns | 13 | 2.40 |
| | Non-contiguous (multiple contigs, non-adjacent loci) | 6 | 1.11 |
| Streptolancidin F | Contiguous | 95 | 100 |
| Streptolancidin G | Contiguous | 252 | 100 |
| Streptolancidin J | Contiguous | 1005 | 99.80 |
| | Non-contiguous (one contig) | 2 | 0.20 |
| Streptolancidin K | Contiguous | 2 | 100 |
| Streptolassin | Contiguous | 48 | 100 |
| Streptosactin | Contiguous | 1 | 100 |

| Kenya | | | |
|---|---|---|---|
| **Bacteriocin cluster** | **Category** | **Frequency** | **% of total bacteriocin clusters** |
| Cib | Contiguous | 3159 | 100 |
| Streptococcin A | Contiguous | 2559 | 99.88 |
| | Non-contiguous (multiple contigs, non-adjacent loci) | 3 | 0.12 |
| Streptococcin B | Contiguous | 3157 | 99.97 |
| | EOC | 1 | 0.03 |
| Streptococcin C | Contiguous | 3152 | 99.78 |
| | EOC | 7 | 0.22 |
| Streptococcin D | Contiguous | 85 | 100 |
| Streptococcin E | Contiguous | 3099 | 98.23 |
| | Non-contiguous (multiple contigs, non-adjacent loci) | 26 | 0.82 |
| | EOC | 16 | 0.51 |
| | Non-contiguous (one contig) | 8 | 0.25 |
| | Non-contiguous (multiple contigs, not EOC-adjacent) | 6 | 0.19 |
| Streptocyclicin | Contiguous | 1516 | 99.48 |
| | EOC | 7 | 0.46 |
| | Non-contiguous (multiple contigs, not EOC-adjacent) | 1 | 0.07 |
| Streptolancidin A | Contiguous | 8 | 100 |
| Streptolancidin B | Contiguous | 338 | 100 |
| Streptolancidin C | Contiguous | 1767 | 97.68 |
| | EOC | 38 | 2.10 |
| | Non-contiguous (multiple contigs, non-adjacent loci) | 3 | 0.17 |
| | Non-contiguous (multiple contigs, not EOC-adjacent) | 1 | 0.06 |
| Streptolancidin D | Contiguous | 850 | 99.65 |
| | Non-contiguous (multiple contigs, non-adjacent loci) | 1 | 0.12 |
| | EOC | 1 | 0.12 |
| | Non-contiguous (multiple contigs, not EOC-adjacent) | 1 | 0.12 |
| Streptolancidin E | Contiguous | 518 | 97.37 |
| | EOC | 8 | 1.50 |

| | | | |
|---|---|---|---|
| | Non-contiguous (multiple contigs, not EOC-adjacent) | 4 | 0.75 |
| | Non-contiguous (multiple contigs, non-adjacent loci) | 1 | 0.19 |
| | Contiguous with Ns | 1 | 0.19 |
| Streptolancidin F | Contiguous | 25 | 100 |
| Streptolancidin G | Contiguous | 272 | 100 |
| Streptolancidin J | Contiguous | 1691 | 99.12 |
| | EOC | 6 | 0.35 |
| | Contiguous with Ns | 5 | 0.29 |
| | Non-contiguous (multiple contigs, not EOC-adjacent) | 3 | 0.18 |
| | Non-contiguous (multiple contigs, non-adjacent loci) | 1 | 0.06 |
| Streptolancidin K | Contiguous | 2 | 100 |
| Streptolassin | Contiguous | 77 | 100 |

Note: Bacteriocin clusters were categorised according to the proximity of the constituent genes to one another. Any clusters with an intergenic region >2.5kbp were categorised as non-contiguous. Bacteriocin clusters with genes on multiple contigs were categorised as 'end of contig' (EOC) if the genes were found within 2.5kbp of each other and the end of the contig, otherwise the clusters were categorised as non-contiguous (ie present on multiple contigs). Rows in grey represent non-contiguous clusters, which were excluded from further analysis.

**Supplementary Table 6: Streptolancidin clusters present in significantly different frequencies among Icelandic and Kenyan pneumococci, stratified by clonal complex (CC).**

| Number of pneumococci harbouring each streptolancidin cluster n (% of CC representatives in each dataset with the bacteriocin) | | |
|---|---|---|
| **Streptolancidin A** | | |
| **CC** | **Iceland** | **Kenya** |
| CC138/176 | 122 (100) | 1 (0.8) |
| CC448 | 29 (100) | 2 (100) |
| CC802 | 0 | 5 (100) |
| CC338 | 1 (12.5) | 0 |
| **Streptolancidin B** | | |
| **CC** | **Iceland** | **Kenya** |
| CC702 | 0 | 57 (98.3) |
| CC499 | 0 | 55 (100) |
| CC5902 | 0 | 32 (13.4) |
| Sing11162 | 0 | 23 (100) |
| CC347 | 0 | 18 (29.0) |
| CC5250/5947/15006 | 0 | 18 (100) |
| CC703 | 0 | 16 (100) |
| CC385 | 0 | 13 (41.9) |
| CC1264 | 0 | 11 (100) |
| CC6446/14764 | 0 | 11 (100) |
| Other CCs | 2 (100) | 62 (34.6) |
| Other Singletons | 0 | 22 (100) |
| **Streptolancidin C** | | |
| **CC** | **Iceland** | **Kenya** |
| CC236/271/320 | 293 (100) | 4 (100) |
| CC138/176 | 122 (100) | 133 (100) |
| CC5902 | 0 | 239 (100) |
| CC217 | 0 | 223 (100) |
| CC5339 | 0 | 138 (97.2) |
| CC156/162 | 0 | 131 (100) |
| CC180 | 107 (100) | 6 (100) |
| CC852 | 0 | 78 (100) |
| CC289 | 0 | 69 (100) |
| CC499 | 0 | 53 (96.4) |
| Other CCs | 262 (52.6) | 652 (71.8) |
| Other Singletons | 14 (100) | 79 (92.9) |
| **Streptolancidin D** | | |
| **CC** | **Iceland** | **Kenya** |
| CC701 | 0 | 161 (98.8) |
| CC5339 | 0 | 139 (97.9) |

| | | |
|---|---|---|
| CC991 | 0 | 104 (100) |
| CC5902 | 0 | 83 (34.7) |
| CC439 | 81 (37.3) | 0 |
| CC854 | 0 | 57 (100) |
| CC706 | 0 | 37 (100) |
| CC15 | 36 (100) | 0 |
| CC14774 | 0 | 23 (100) |
| Sing11162 | 0 | 23 (100) |
| Other CCs | 55 (25.0) | 190 (42.6) |
| Other Singletons | 12 (100) | 34 (100) |
| **Streptolancidin E** | | |
| **CC** | **Iceland** | **Kenya** |
| CC439 | 217 (100) | 0 |
| CC199 | 174 (97.2) | 0 |
| CC1146 | 0 | 99 (71.2) |
| CC230 | 3 (100) | 88 (95.7) |
| CC5258 | 0 | 76 (98.7) |
| CC1381 | 0 | 49 (100) |
| CC344 | 37 (100) | 1 (100) |
| CC705/14790 | 0 | 38 (100) |
| CC448 | 29 (100) | 2 (100) |
| CC138/176 | 0 | 22 (16.5) |
| Other CCs | 59 (43.4) | 115 (29.0) |
| Other Singletons | 2 (100) | 37 (100) |
| **Streptolancidin F** | | |
| **CC** | **Iceland** | **Kenya** |
| CC344 | 33 (89.2) | 1 (100) |
| CC100 | 25 (100) | 0 |
| CC191 | 16 (100) | 0 |
| CC5560/6090/6103 | 0 | 14 (100) |
| CC433 | 9 (14.8) | 0 |
| CC5292 | 0 | 7 (100) |
| CC717 | 4 (100) | 0 |
| CC97 | 3 (3.4) | 0 |
| CC346 | 2 (100) | 0 |
| CC113 | 2 (9.5) | 0 |
| Other CCs | 0 | 2 (2.2) |
| Other Singletons | 1 (100) | 1 (100) |
| **Streptolancidin G** | | |
| **CC** | **Iceland** | **Kenya** |
| CC1146 | 0 | 134 (96.4) |
| CC852 | 0 | 78 (100) |

| | | |
|---|---|---|
| CC433 | 61 (100) | 0 |
| CC392 | 47 (100) | 0 |
| CC5329 | 0 | 37 (97.4) |
| CC393 | 20 (100) | 9 (100) |
| CC66 | 18 (94.7) | 0 |
| CC30 | 17 (28.3) | 0 |
| CC2755 | 16 (100) | 0 |
| CC315 | 13 (86.7) | 0 |
| Other CCs | 48 (63.2) | 12 (4.0) |
| Other Singletons | 12 (100) | 2 (100) |

Note: The 10 CCs with the biggest contribution to the frequency of each streptolancidin are shown. Other CCs were pooled to the 'Other' categories.

**Supplementary Table 7: Association of bacteriocin clusters with pneumococcal serotype.**

| Streptococcin A | | | |
|---|---|---|---|
| Iceland | | Kenya | |
| Significant in IPD, OM, LRTI | | Significant in IPD | |
| Serotype | n (% of all pneumococci with that serotype) | Serotype | n (% of all pneumococci with that serotype) |
| 19F | 322 (97.3) | 1 | 223 (99.6) |
| 6A | 163 (100) | 19F | 211 (92.5) |
| 23F | 151 (83.9) | 6A | 194 (94.2) |
| 6B | 121 (99.2) | 19A | 153 (100) |
| 3 | 107 (98.2) | 35B | 139 (100) |
| 11A | 93 (100) | 15A | 136 (100) |
| 14 | 81 (90.0) | 15BC | 130 (89.0) |
| 22F | 61 (100) | 11A | 116 (99.1) |
| 23B | 47 (100) | 13 | 97 (99.0) |
| 23A | 45 (88.2) | 14 | 95 (72.5) |
| Other serotypes | 346 (61.3) | Other serotypes | 1065 (68.9) |
| **Streptococcin D** | | | |
| Iceland | | Kenya | |
| Not significant | | Significant in IPD | |
| Serotype | n (% of all pneumococci with that serotype) | Serotype | n (% of all pneumococci with that serotype) |
| - | - | 14 | 70 (53.4) |
| - | - | nontypable | 15 (46.9) |
| **Streptococcin E** | | | |
| Iceland | | Kenya | |
| Significant in IPD, OM, LRTI | | Significant in IPD | |
| Serotype | n (% of all pneumococci with that serotype) | Serotype | n (% of all pneumococci with that serotype) |
| 19F | 328 (99.1) | 19F | 228 (100) |
| 23F | 180 (100) | 1 | 224 (100) |
| 6A | 163 (100) | 6A | 206 (100) |
| 19A | 145 (100) | 19A | 153 (100) |
| 6B | 122 (100) | 15BC | 146 (100) |
| 3 | 109 (100) | 35B | 139 (100) |
| 11A | 93 (100) | 15A | 136 (100) |
| 15BC | 93 (100) | 14 | 131 (100) |
| 14 | 90 (100) | 6E(6Bii) | 131 (100) |

| | | | |
|---|---|---|---|
| 22F | 61 (100) | 23F | 119 (100) |
| Other serotypes | 456 (86.9) | Other serotypes | 1502 (97.2) |
| **Streptocyclicin** | | | |
| **Iceland** | | **Kenya** | |
| **Significant in carriage** | | **Significant in carriage** | |
| **Serotype** | **n (% of all pneumococci with that serotype)** | **Serotype** | **n (% of all pneumococci with that serotype)** |
| 23F | 174 (96.7) | 19A | 147 (96.1) |
| 19A | 127 (87.6) | 15A | 126 (92.6) |
| 15BC | 93 (100) | 6A | 125 (60.7) |
| 6A | 69 (42.3) | 13 | 98 (100) |
| 14 | 66 (73.3) | 11A | 94 (80.3) |
| 23A | 50 (98.0) | 16F | 93 (96.9) |
| 23B | 43 (91.5) | 23B | 90 (100) |
| nontypable | 36 (51.4) | 34 | 85 (95.5) |
| 9V | 32 (100) | 10A | 82 (100) |
| 16F | 26 (100) | 5 | 69 (100) |
| Other serotypes | 146 (25.0) | Other serotypes | 514 (29.8) |
| **Streptolancidin A** | | | |
| **Iceland** | | **Kenya** | |
| **Significant in carriage** | | **Not significant** | |
| **Serotype** | **n (% of all pneumococci with that serotype)** | **Serotype** | **n (% of all pneumococci with that serotype)** |
| 6B | 120 (98.4) | - | - |
| nontypable | 29 (41.4) | - | - |
| 6A | 3 (1.8) | - | - |
| **Streptolancidin B** | | | |
| **Iceland** | | **Kenya** | |
| **Not significant** | | **Significant in carriage** | |
| **Serotype** | **n (% of all pneumococci with that serotype)** | **Serotype** | **n (% of all pneumococci with that serotype)** |
| - | - | 6A | 64 (31.1) |
| - | - | 20 | 53 (98.1) |
| - | - | 15BC | 47 (32.2) |
| - | - | 11A | 33 (28.2) |
| - | - | 16F | 32 (33.3) |
| - | - | 19F | 23 (10.1) |
| - | - | 6E(6Bii) | 20 (15.3) |

| | | | |
|---|---|---|---|
| - | - | 15A | 12 (8.8) |
| - | - | 24F | 12 (100) |
| - | - | 6C | 10 (100) |
| - | - | Other serotypes | 32 (5.1) |
| **Streptolancidin C** | | | |
| **Iceland** | | **Kenya** | |
| **Significant in OM, LRTI** | | **Significant in IPD** | |
| **Serotype** | **n (% of all pneumococci with that serotype)** | **Serotype** | **n (% of all pneumococci with that serotype)** |
| 19F | 325 (98.2) | 1 | 224 (100) |
| 6B | 122 (100) | 19F | 180 (78.9) |
| 3 | 109 (100) | 19A | 149 (97.4) |
| 23A | 40 (78.4) | 6A | 119 (57.8) |
| 6A | 29 (17.8) | 23F | 119 (100) |
| nontypable | 29 (41.4) | 11A | 112 (95.7) |
| 6E | 24 (92.3) | 15BC | 88 (60.3) |
| 14 | 21 (23.3) | 23B | 87 (96.7) |
| 38 | 20 (100) | 10A | 81 (98.8) |
| 7F | 16 (100) | 5 | 69 (100) |
| Other Serotypes | 63 (10.8) | Other serotypes | 577 (37.9) |
| **Streptolancidin D** | | | |
| **Iceland** | | **Kenya** | |
| **Significant in OM** | | **Significant in carriage** | |
| **Serotype** | **n (% of all pneumococci with that serotype)** | **Serotype** | **n (% of all pneumococci with that serotype)** |
| 23F | 83 (46.1) | 19F | 150 (65.8) |
| 6A | 22 (13.5) | 15A | 124 (91.2) |
| 14 | 21 (23.3) | 15BC | 113 (77.4) |
| 19F | 17 (5.1) | 13 | 97 (99.0) |
| 35B | 16 (48.5) | 11A | 75 (64.1) |
| 19A | 12 (8.3) | 6E(6Bii) | 57 (43.5) |
| 6C | 8 (27.6) | 9V | 51 (81.0) |
| 18C | 2 (9.1) | 21 | 25 (40.3) |
| 31 | 1 (33.3) | 6A | 22 (10.7) |
| 6E | 1 (3.8) | 6B | 21 (80.8) |
| Other serotypes | 1 (12.5) | Other serotypes | 116 (13.5) |
| **Streptolancidin E** | | | |
| **Iceland** | | **Kenya** | |
| **Significant in carriage** | | **Significant in carriage** | |

| Serotype | n (% of all pneumococci with that serotype) | Serotype | n (% of all pneumococci with that serotype) |
|---|---|---|---|
| 23F | 127 (70.6) | 35B | 96 (69.1) |
| 19A | 126 (86.9) | 34 | 76 (85.4) |
| nontypable | 67 (95.7) | 16F | 57 (59.4) |
| 15BC | 54 (58.1) | 3 | 56 (62.2) |
| 23A | 50 (98.0) | 18C | 48 (100) |
| 23B | 42 (89.4) | 14 | 34 (26.0) |
| 18C | 17 (77.3) | 17F | 20 (80.0) |
| 9N | 14 (77.8) | 21 | 20 (32.3) |
| 21 | 4 (13.8) | 35F | 17 (94.4) |
| 1 | 3 (60.0) | 15BC | 14 (9.6) |
| Other serotypes | 17 (2.6) | Other serotypes | 89 (9.4) |
| **Streptolancidin F** | | | |
| **Iceland** | | **Kenya** | |
| **Significant in IPD, carriage (vs OM)** | | **Not significant** | |
| **Serotype** | **n (% of all pneumococci with that serotype)** | **Serotype** | **n (% of all pneumococci with that serotype)** |
| NT | 34 (48.6) | - | - |
| 33F | 29 (100) | - | - |
| 7F | 16 (100) | - | - |
| 22F | 9 (14.8) | - | - |
| 10A | 3 (60.0) | - | - |
| 19A | 2 (1.4) | - | - |
| 18C | 2 (9.1) | - | - |
| **Streptolancidin G** | | | |
| **Iceland** | | **Kenya** | |
| **Significant in IPD, carriage (vs OM)** | | **Significant in carriage** | |
| **Serotype** | **n (% of all pneumococci with that serotype)** | **Serotype** | **n (% of all pneumococci with that serotype)** |
| 22F | 60 (98.4) | 35B | 134 (96.4) |
| 23F | 48 (26.7) | 10A | 82 (100) |
| 35B | 30 (90.9) | 29 | 19 (67.9) |
| 38 | 20 (100) | 6A | 12 (5.8) |
| 6C | 20 (69.0) | 38 | 8 (34.8) |
| 9N | 18 (100) | 6E(6Bii) | 4 (3.1) |
| 19F | 17 (5.1) | 10F | 3 (33.3) |

| | | | |
|---|---|---|---|
| 6A | 14 (8.6) | 15BC | 2 (1.4) |
| 19A | 12 (8.3) | 21 | 2 (3.2) |
| 4 | 7 (100) | 34 | 1 (1.1) |
| Other serotypes | 6 (10.3) | Other serotypes | 5 (1.5) |
| **Streptolancidin J** | | | |
| **Iceland** | | **Kenya** | |
| **Significant in carriage** | | **Not significant** | |
| **Serotype** | **n (% of all pneumococci with that serotype)** | **Serotype** | **n (% of all pneumococci with that serotype)** |
| 6A | 147 (90.2) | - | - |
| 19A | 139 (95.9) | - | - |
| 6B | 122 (100) | - | - |
| 3 | 107 (98.2) | - | - |
| 14 | 69 (76.7) | - | - |
| 15BC | 62 (66.7) | - | - |
| 22F | 61 (100) | - | - |
| 23F | 50 (27.8) | - | - |
| 19F | 34 (10.3) | - | - |
| 9V | 31 (96.9) | - | - |
| Other serotypes | 183 (35.6) | - | - |
| **Streptolassin** | | | |
| **Iceland** | | **Kenya** | |
| **Significant in carriage (v OM)** | | **Significant in IPD** | |
| **Serotype** | **n (% of all pneumococci with that serotype)** | **Serotype** | **n (% of all pneumococci with that serotype)** |
| 23F | 48 (26.7) | 5 | 69 (100) |
| - | - | 37 | 3 (100) |
| - | - | 7F | 1 (50.0) |
| - | - | 1 | 1 (0.4) |
| - | - | 38 | 1 (4.3) |
| - | - | 6A | 1 (0.5) |
| - | - | 8 | 1 (4.8) |

Note: Up to 10 of the most common serotypes associated with each bacteriocin are listed separately, and the remainder were pooled as "Other". Bacteriocins that did not exhibit significantly altered prevalence in any subset of the data were excluded from this table.

**Supplementary Table 8: Bacteriocin clusters in the Icelandic dataset.**

| Number of pneumococci harbouring each bacteriocin cluster, stratified by CC<br>n (% of CC representatives in each subset with the bacteriocin) | | | | | | |
|---|---|---|---|---|---|---|
| **Streptococcin A** | | | | | | |
| **CC** | **Pre-PCV** | **Post-PCV** | **Carriage** | **IPD** | **LRTI** | **OM** |
| CC236/271/320 | 201 (97.6) | 85 (97.7) | 53 (100) | 6 (100) | 72 (93.5) | 155 (98.7) |
| CC439 | 86 (80.4) | 98 (89.1) | 106 (83.5) | 16 (94.1) | 21 (95.5) | 41 (80.4) |
| CC138/176 | 79 (100) | 42 (97.7) | 87 (100) | 5 (100) | 11 (91.7) | 18 (100) |
| CC180 | 64 (100) | 42 (97.7) | 55 (100) | 9 (100) | 21 (100) | 21 (95.5) |
| CC62 | 37 (97.4) | 56 (100) | 62 (100) | 5 (100) | 13 (92.9) | 13 (100) |
| CC490 | 40 (100) | 34 (100) | 46 (100) | 5 (100) | 9 (100) | 14 (100) |
| CC433 | 13 (100) | 48 (100) | 32 (100) | 13 (100) | 11 (100) | 5 (100) |
| CC30 | 34 (100) | 26 (100) | 40 (100) | 2 (100) | 10 (100) | 8 (100) |
| CC97 | 30 (88.2) | 30 (56.6) | 32 (66.7) | 4 (66.7) | 4 (44.4) | 20 (83.3) |
| CC124 | 36 (81.8) | 17 (94.4) | 23 (79.3) | 12 (100) | 5 (71.4) | 13 (92.9) |
| Other CCs | 213 (91.8) | 211 (96.8) | 200 (91.7) | 75 (93.8) | 65 (98.5) | 84 (97.7) |
| Other Singletons | 4 (100) | 11 (100) | 10 (100) | 1 (100) | 1 (100) | 3 (100) |
| **Streptococcin E** | | | | | | |
| **CC** | **Pre-PCV** | **Post-PCV** | **Carriage** | **IPD** | **LRTI** | **OM** |
| CC236/271/320 | 203 (98.5) | 87 (100) | 53 (100) | 6 (100) | 75 (97.4) | 156 (99.4) |
| CC439 | 107 (100) | 110 (100) | 127 (100) | 17 (100) | 22 (100) | 51 (100) |
| CC199 | 99 (100) | 80 (100) | 110 (100) | 13 (100) | 11 (100) | 45 (100) |
| CC138/176 | 79 (100) | 43 (100) | 87 (100) | 5 (100) | 12 (100) | 18 (100) |
| CC180 | 64 (100) | 43 (100) | 55 (100) | 9 (100) | 21 (100) | 22 (100) |
| CC62 | 38 (100) | 56 (100) | 62 (100) | 5 (100) | 14 (100) | 13 (100) |
| CC97 | 34 (100) | 53 (100) | 48 (100) | 6 (100) | 9 (100) | 24 (100) |
| CC490 | 40 (100) | 34 (100) | 46 (100) | 5 (100) | 9 (100) | 14 (100) |
| CC124 | 44 (100) | 18 (100) | 29 (100) | 12 (100) | 7 (100) | 14 (100) |
| CC433 | 13 (100) | 48 (100) | 32 (100) | 13 (100) | 11 (100) | 5 (100) |
| Other CCs | 279 (99.6) | 253 (100) | 262 (99.6) | 91 (100) | 83 (100) | 96 (100) |
| Other Singletons | 4 (100) | 11 (100) | 10 (100) | 1 (100) | 1 (100) | 3 (100) |
| **Streptocyclicin** | | | | | | |
| **CC** | **Pre-PCV** | **Post-PCV** | **Carriage** | **IPD** | **LRTI** | **OM** |
| CC439 | 107 (100) | 110 (100) | 127 (100) | 17 (100) | 22 (100) | 51 (100) |
| CC199 | 99 (100) | 80 (100) | 110 (100) | 13 (100) | 11 (100) | 45 (100) |
| CC97 | 34 (100) | 53 (100) | 48 (100) | 6 (100) | 9 (100) | 24 (100) |
| CC124 | 44 (100) | 18 (100) | 29 (100) | 12 (100) | 7 (100) | 14 (100) |
| CC392 | 24 (88.9) | 19 (95.0) | 31 (96.9) | 1 (33.3) | 5 (100) | 6 (85.7) |
| CC156/162 | 36 (92.3) | 7 (100) | 14 (82.4) | 11 (100) | 10 (100) | 8 (100) |
| CC30 | 24 (70.6) | 19 (73.1) | 32 (80.0) | 1 (50.0) | 6 (60.0) | 4 (50.0) |
| CC1262 | 6 (100) | 29 (100) | 21 (100) | 2 (100) | 7 (100) | 5 (100) |
| CC344 | 14 (87.5) | 21 (100) | 31 (93.9) | 0 | 4 (100) | 0 |

| | | | | | | |
|---|---|---|---|---|---|---|
| CC193 | 6 (100) | 22 (100) | 16 (100) | 2 (100) | 5 (100) | 5 (100) |
| Other CCs | 33 (71.7) | 55 (53.9) | 50 (61.7) | 16 (55.2) | 11 (50.0) | 11 (68.8) |
| Other Singletons | 0 | 2 (100) | 1 (100) | 0 | 0 | 1 (100) |
| **Streptolancidin A** | | | | | | |
| **CC** | **Pre-PCV** | **Post-PCV** | **Carriage** | **IPD** | **LRTI** | **OM** |
| CC138/176 | 79 (100) | 43 (100) | 87 (100) | 5 (100) | 12 (100) | 18 (100) |
| CC448 | 15 (100) | 14 (100) | 27 (100) | 0 | 2 (100) | 0 |
| CC338 | 1 (20.0) | 0 | 0 | 1 (50.0) | 0 | 0 |
| **Streptolancidin C** | | | | | | |
| **CC** | **Pre-PCV** | **Post-PCV** | **Carriage** | **IPD** | **LRTI** | **OM** |
| CC236/271/320 | 206 (100) | 87 (100) | 53 (100) | 6 (100) | 77 (100) | 157 (100) |
| CC138/176 | 79 (100) | 43 (100) | 87 (100) | 5 (100) | 12 (100) | 18 (100) |
| CC180 | 64 (100) | 43 (100) | 55 (100) | 9 (100) | 21 (100) | 22 (100) |
| CC439 | 10 (9.3) | 32 (29.1) | 31 (24.4) | 2 (11.8) | 3 (13.6) | 6 (11.8) |
| CC15 | 32 (100) | 4 (100) | 14 (100) | 9 (100) | 5 (100) | 8 (100) |
| CC30 | 15 (44.1) | 19 (73.1) | 20 (50.0) | 2 (100) | 6 (60.0) | 6 (75.0) |
| CC448 | 15 (100) | 14 (100) | 27 (100) | 0 | 2 (100) | 0 |
| CC90 | 15 (100) | 7 (100) | 8 (100) | 1 (100) | 7 (100) | 6 (100) |
| CC393 | 16 (100) | 4 (100) | 16 (100) | 2 (100) | 0 | 2 (100) |
| CC191 | 11 (100) | 5 (100) | 0 | 14 (100) | 1 (100) | 1 (100) |
| Other CCs | 29 (100) | 34 (100) | 26 (100) | 11 (100) | 13 (100) | 13 (100) |
| Other Singletons | 3 (100) | 11 (100) | 10 (100) | 0 | 1 (100) | 3 (100) |
| **Streptolancidin D** | | | | | | |
| **CC** | **Pre-PCV** | **Post-PCV** | **Carriage** | **IPD** | **LRTI** | **OM** |
| CC439 | 51 (47.7) | 30 (27.3) | 34 (26.8) | 9 (52.9) | 10 (45.5) | 28 (54.9) |
| CC15 | 32 (100) | 4 (100) | 14 (100) | 9 (100) | 5 (100) | 8 (100) |
| CC30 | 5 (14.7) | 12 (46.2) | 12 (30.0) | 1 (50.0) | 2 (20.0) | 2 (25.0) |
| CC2755 | 8 (100) | 8 (100) | 4 (100) | 3 (100) | 5 (100) | 4 (100) |
| CC177 | 6 (100) | 8 (100) | 5 (100) | 0 | 3 (100) | 6 (100) |
| Sing1801 | 2 (100) | 10 (100) | 9 (100) | 0 | 1 (100) | 2 (100) |
| CC473 | 1 (100) | 1 (100) | 1 (100) | 0 | 0 | 1 (100) |
| CC102 | 2 (100) | 0 | 2 (100) | 0 | 0 | 0 |
| CC338 | 2 (40.0) | 0 | 0 | 0 | 1 (50.0) | 1 (50.0) |
| CC1766 | 1 (100) | 0 | 0 | 0 | 1 (100) | 0 |
| Other CCs | 1 (100) | 0 | 0 | 0 | 0 | 1 (25.0) |
| Other Singletons | 0 | 0 | 0 | 0 | 0 | 0 |
| **Streptolancidin E** | | | | | | |
| **CC** | **Pre-PCV** | **Post-PCV** | **Carriage** | **IPD** | **LRTI** | **OM** |
| CC439 | 107 (100) | 110 (100) | 127 (100) | 17 (100) | 22 (100) | 51 (100) |
| CC199 | 97 (98.0) | 77 (96.2) | 108 (98.2) | 13 (100) | 10 (90.9) | 43 (95.6) |
| CC344 | 16 (100) | 21 (100) | 33 (100) | 0 | 4 (100) | 0 |
| CC448 | 15 (100) | 14 (100) | 27 (100) | 0 | 2 (100) | 0 |

| | | | | | | |
|---|---|---|---|---|---|---|
| CC113 | 12 (85.7) | 6 (85.7) | 12 (80.0) | 2 (100) | 2 (100) | 2 (100) |
| CC66 | 8 (88.9) | 7 (70.0) | 9 (90.0) | 4 (66.7) | 1 (50.0) | 1 (100) |
| CC3017 | 6 (100) | 2 (100) | 2 (100) | 3 (100) | 3 (100) | 0 |
| CC432 | 0 | 4 (66.7) | 3 (60.0) | 0 | 0 | 1 (100) |
| CC306 | 1 (50.0) | 2 (66.7) | 0 | 3 (60.0) | 0 | 0 |
| CC230 | 1 (100) | 2 (100) | 0 | 1 (100) | 0 | 2 (100) |
| Other CCs | 7 (17.5) | 4 (13.8) | 3 (7.3) | 3 (60.0) | 5 (33.3) | 0 |
| Other Singletons | 1 (100) | 1 (100) | 1 (100) | 1 (100) | 0 | 0 |
| **Streptolancidin F** | | | | | | |
| **CC** | **Pre-PCV** | **Post-PCV** | **Carriage** | **IPD** | **LRTI** | **OM** |
| CC344 | 14 (87.5) | 19 (90.5) | 29 (87.9) | 0 | 4 (100) | 0 |
| CC100 | 15 (100) | 10 (100) | 10 (100) | 7 (100) | 4 (100) | 4 (100) |
| CC191 | 11 (100) | 5 (100) | 0 | 14 (100) | 1 (100) | 1 (100) |
| CC433 | 7 (53.8) | 2 (4.2) | 2 (6.2) | 2 (15.4) | 5 (45.5) | 0 |
| CC717 | 1 (100) | 3 (100) | 1 (100) | 0 | 0 | 3 (100) |
| CC97 | 2 (5.9) | 1 (1.9) | 3 (6.2) | 0 | 0 | 0 |
| CC113 | 2 (14.3) | 0 | 2 (13.3) | 0 | 0 | 0 |
| CC346 | 0 | 2 (100) | 1 (100) | 0 | 0 | 1 (100) |
| Sing10346 | 0 | 1 (100) | 1 (100) | 0 | 0 | 0 |
| **Streptolancidin G** | | | | | | |
| **CC** | **Pre-PCV** | **Post-PCV** | **Carriage** | **IPD** | **LRTI** | **OM** |
| CC433 | 13 (100) | 48 (100) | 32 (100) | 13 (100) | 11 (100) | 5 (100) |
| CC392 | 27 (100) | 20 (100) | 32 (100) | 3 (100) | 5 (100) | 7 (100) |
| CC393 | 16 (100) | 4 (100) | 16 (100) | 2 (100) | 0 | 2 (100) |
| CC66 | 9 (100) | 9 (90.0) | 10 (100) | 5 (83.3) | 2 (100) | 1 (100) |
| CC30 | 10 (29.4) | 7 (26.9) | 8 (20.0) | 1 (50.0) | 4 (40.0) | 4 (50.0) |
| CC2755 | 8 (100) | 8 (100) | 4 (100) | 3 (100) | 5 (100) | 4 (100) |
| CC315 | 4 (66.7) | 9 (100) | 4 (80.0) | 1 (100) | 2 (100) | 6 (85.7) |
| CC15 | 12 (37.5) | 1 (25.0) | 8 (57.1) | 0 | 3 (60.0) | 2 (25.0) |
| CC198 | 0 | 13 (100) | 11 (100) | 0 | 1 (100) | 1 (100) |
| Sing1801 | 2 (100) | 10 (100) | 9 (100) | 0 | 1 (100) | 2 (100) |
| Other CCs | 8 (100) | 14 (73.7) | 7 (63.6) | 9 (100) | 2 (100) | 4 (80.0) |
| Other Singletons | 0 | 0 | 0 | 0 | 0 | 0 |
| **Streptolancidin J** | | | | | | |
| **CC** | **Pre-PCV** | **Post-PCV** | **Carriage** | **IPD** | **LRTI** | **OM** |
| CC199 | 86 (86.9) | 67 (83.8) | 97 (88.2) | 11 (84.6) | 7 (63.6) | 38 (84.4) |
| CC138/176 | 79 (100) | 43 (100) | 87 (100) | 5 (100) | 12 (100) | 18 (100) |
| CC180 | 64 (100) | 43 (100) | 55 (100) | 9 (100) | 21 (100) | 22 (100) |
| CC97 | 34 (100) | 52 (98.1) | 48 (100) | 6 (100) | 8 (88.9) | 24 (100) |
| CC490 | 39 (97.5) | 32 (94.1) | 44 (95.7) | 5 (100) | 9 (100) | 13 (92.9) |
| CC124 | 44 (100) | 18 (100) | 29 (100) | 12 (100) | 7 (100) | 14 (100) |
| CC433 | 13 (100) | 48 (100) | 32 (100) | 13 (100) | 11 (100) | 5 (100) |

| | | | | | | |
|---|---|---|---|---|---|---|
| CC30 | 34 (100) | 26 (100) | 40 (100) | 2 (100) | 10 (100) | 8 (100) |
| CC392 | 27 (100) | 20 (100) | 32 (100) | 3 (100) | 5 (100) | 7 (100) |
| CC156/162 | 38 (97.4) | 7 (100) | 17 (100) | 10 (90.9) | 10 (100) | 8 (100) |
| Other CCs | 66 (60.6) | 113 (57.4) | 91 (52.3) | 24 (68.6) | 29 (63.0) | 35 (68.6) |
| Other Singletons | 2 (100) | 10 (100) | 9 (100) | 0 | 1 (100) | 2 (100) |
| **Streptolassin** | | | | | | |
| **CC** | **Pre-PCV** | **Post-PCV** | **Carriage** | **IPD** | **LRTI** | **OM** |
| CC392 | 27 (100) | 20 (100) | 32 (100) | 3 (100) | 5 (100) | 7 (100) |
| CC433 | 0 | 1 (2.1) | 1 (3.1) | 0 | 0 | 0 |

Note: The 10 most common clonal complexes in which the bacteriocin cluster was found are listed separately, and the remainder were pooled as 'Other'. Only bacteriocin clusters with significant differences in prevalence in Figure 2B and 2C are included in this table. IPD, invasive pneumococcal disease, LRTI, lower respiratory tract infection, OM, otitis media.

**Supplementary Table 9: Bacteriocin clusters in the Kenyan dataset.**

| **Number of pneumococci harbouring each bacteriocin cluster, stratified by CC n (% of CC representatives in each subset with the bacteriocin)** | | | | |
|---|---|---|---|---|
| **Streptococcin A** | | | | |
| **CC** | **Pre-PCV** | **Post-PCV** | **Carriage** | **IPD** |
| CC5902 | 101 (100) | 135 (97.8) | 219 (98.6) | 17 (100) |
| CC217 | 199 (99.5) | 23 (100) | 16 (100) | 206 (99.5) |
| CC701 | 67 (93.1) | 80 (87.9) | 130 (89.7) | 17 (94.4) |
| CC1146 | 53 (100) | 86 (100) | 126 (100) | 13 (100) |
| CC5339 | 106 (97.2) | 33 (100) | 122 (97.6) | 17 (100) |
| CC156/162 | 36 (100) | 95 (100) | 97 (100) | 34 (100) |
| CC138/176 | 66 (98.5) | 64 (97.0) | 97 (98.0) | 33 (97.1) |
| CC991 | 24 (100) | 80 (100) | 95 (100) | 9 (100) |
| CC852 | 26 (96.3) | 51 (100) | 66 (98.5) | 11 (100) |
| CC63 | 56 (100) | 14 (100) | 37 (100) | 33 (100) |
| Other CCs | 576 (81.5) | 472 (82.8) | 801 (81.2) | 247 (85.2) |
| Other Singletons | 42 (100) | 74 (98.7) | 100 (99.0) | 16 (100) |
| **Streptococcin D** | | | | |
| **CC** | **Pre-PCV** | **Post-PCV** | **Carriage** | **IPD** |
| CC63 | 56 (100) | 14 (100) | 37 (100) | 33 (100) |
| CC13215 | 0 | 14 (100) | 14 (100) | 0 |
| Sing14766 | 1 (100) | 0 | 0 | 1 (100) |
| **Streptococcin E** | | | | |
| **CC** | **Pre-PCV** | **Post-PCV** | **Carriage** | **IPD** |
| CC5902 | 101 (100) | 138 (100) | 222 (100) | 17 (100) |
| CC217 | 200 (100) | 23 (100) | 16 (100) | 207 (100) |
| CC701 | 72 (100) | 91 (100) | 145 (100) | 18 (100) |

| | | | | |
|---|---|---|---|---|
| CC5339 | 109 (100) | 33 (100) | 125 (100) | 17 (100) |
| CC1146 | 53 (100) | 86 (100) | 126 (100) | 13 (100) |
| CC138/176 | 67 (100) | 66 (100) | 99 (100) | 34 (100) |
| CC156/162 | 36 (100) | 95 (100) | 97 (100) | 34 (100) |
| CC991 | 24 (100) | 80 (100) | 95 (100) | 9 (100) |
| CC230 | 50 (100) | 42 (100) | 60 (100) | 32 (100) |
| CC852 | 27 (100) | 51 (100) | 67 (100) | 11 (100) |
| Other CCs | 857 (98.6) | 671 (96.1) | 1166 (96.8) | 362 (99.7) |
| Other Singletons | 49 (100) | 94 (100) | 127 (100) | 16 (100) |
| **Streptocyclicin** | | | | |
| **CC** | **Pre-PCV** | **Post-PCV** | **Carriage** | **IPD** |
| CC5902 | 78 (77.2) | 89 (64.5) | 152 (68.5) | 15 (88.2) |
| CC701 | 72 (100) | 91 (100) | 145 (100) | 18 (100) |
| CC156/162 | 36 (100) | 95 (100) | 97 (100) | 34 (100) |
| CC991 | 24 (100) | 80 (100) | 95 (100) | 9 (100) |
| CC230 | 48 (96.0) | 41 (97.6) | 58 (96.7) | 31 (96.9) |
| CC852 | 27 (100) | 51 (100) | 67 (100) | 11 (100) |
| CC5258 | 18 (100) | 59 (100) | 72 (100) | 5 (100) |
| CC289 | 64 (100) | 5 (100) | 3 (100) | 66 (100) |
| CC914 | 43 (97.7) | 17 (100) | 44 (97.8) | 16 (100) |
| CC702 | 18 (100) | 40 (100) | 56 (100) | 2 (100) |
| Other CCs | 213 (45.6) | 257 (71.2) | 395 (58.1) | 75 (50.7) |
| Other Singletons | 23 (100) | 34 (100) | 51 (100) | 6 (100) |
| **Streptolancidin B** | | | | |
| **CC** | **Pre-PCV** | **Post-PCV** | **Carriage** | **IPD** |
| CC702 | 17 (94.4) | 40 (100) | 55 (98.2) | 2 (100) |
| CC499 | 36 (100) | 19 (100) | 44 (100) | 11 (100) |
| CC5902 | 16 (15.8) | 16 (11.6) | 32 (14.4) | 0 |
| Sing11162 | 0 | 23 (100) | 20 (100) | 3 (100) |
| CC347 | 16 (28.1) | 2 (40.0) | 12 (25.0) | 6 (42.9) |
| CC5250/5947/15006 | 10 (100) | 8 (100) | 16 (100) | 2 (100) |
| CC703 | 9 (100) | 7 (100) | 14 (100) | 2 (100) |
| CC385 | 10 (41.7) | 3 (42.9) | 6 (37.5) | 7 (46.7) |
| CC1264 | 3 (100) | 8 (100) | 11 (100) | 0 |
| CC6446/14764 | 2 (100) | 9 (100) | 9 (100) | 2 (100) |
| Other CCs | 40 (40.0) | 22 (27.8) | 55 (36.7) | 7 (24.1) |
| Other Singletons | 10 (100) | 12 (100) | 19 (100) | 3 (100) |
| **Streptolancidin C** | | | | |
| **CC** | **Pre-PCV** | **Post-PCV** | **Carriage** | **IPD** |
| CC5902 | 101 (100) | 138 (100) | 222 (100) | 17 (100) |
| CC217 | 200 (100) | 23 (100) | 16 (100) | 207 (100) |
| CC5339 | 106 (97.2) | 32 (97.0) | 121 (96.8) | 17 (100) |

| | | | | |
|---|---|---|---|---|
| CC138/176 | 67 (100) | 66 (100) | 99 (100) | 34 (100) |
| CC156/162 | 36 (100) | 95 (100) | 97 (100) | 34 (100) |
| CC852 | 27 (100) | 51 (100) | 67 (100) | 11 (100) |
| CC289 | 64 (100) | 5 (100) | 3 (100) | 66 (100) |
| CC499 | 36 (100) | 17 (89.5) | 42 (95.5) | 11 (100) |
| CC7689 | 36 (100) | 3 (100) | 30 (100) | 9 (100) |
| CC338 | 16 (100) | 21 (100) | 29 (100) | 8 (100) |
| Other CCs | 285 (66.6) | 301 (72.7) | 485 (70.0) | 101 (67.8) |
| Other Singletons | 24 (88.9) | 55 (94.8) | 70 (92.1) | 9 (100) |
| **Streptolancidin D** | | | | |
| **CC** | **Pre-PCV** | **Post-PCV** | **Carriage** | **IPD** |
| CC701 | 71 (98.6) | 90 (98.9) | 143 (98.6) | 18 (100) |
| CC5339 | 107 (98.2) | 32 (97.0) | 122 (97.6) | 17 (100) |
| CC991 | 24 (100) | 80 (100) | 95 (100) | 9 (100) |
| CC5902 | 45 (44.6) | 38 (27.5) | 75 (33.8) | 8 (47.1) |
| CC854 | 52 (100) | 5 (100) | 38 (100) | 19 (100) |
| CC706 | 30 (100) | 7 (100) | 27 (100) | 10 (100) |
| Sing11162 | 0 | 23 (100) | 20 (100) | 3 (100) |
| CC14774 | 6 (100) | 17 (100) | 21 (100) | 2 (100) |
| CC4368 | 17 (94.4) | 5 (100) | 17 (94.4) | 5 (100) |
| CC5938 | 9 (100) | 9 (90.0) | 16 (94.1) | 2 (100) |
| Other CCs | 68 (43.0) | 82 (46.6) | 131 (51.8) | 19 (23.5) |
| Other Singletons | 14 (100) | 20 (100) | 29 (100) | 5 (100) |
| **Streptolancidin E** | | | | |
| **CC** | **Pre-PCV** | **Post-PCV** | **Carriage** | **IPD** |
| CC1146 | 45 (84.9) | 54 (62.8) | 87 (69.0) | 12 (92.3) |
| CC230 | 48 (96.0) | 40 (95.2) | 57 (95.0) | 31 (96.9) |
| CC5258 | 18 (100) | 58 (98.3) | 71 (98.6) | 5 (100) |
| CC1381 | 41 (100) | 8 (100) | 31 (100) | 18 (100) |
| CC705/14790 | 8 (100) | 30 (100) | 33 (100) | 5 (100) |
| CC138/176 | 5 (7.5) | 17 (25.8) | 17 (17.2) | 5 (14.7) |
| CC5349 | 6 (100) | 11 (100) | 17 (100) | 0 |
| CC14858 | 5 (100) | 11 (73.3) | 14 (77.8) | 2 (100) |
| CC14892 | 1 (16.7) | 13 (81.2) | 14 (63.6) | 0 |
| Sing14868 | 5 (100) | 9 (100) | 12 (100) | 2 (100) |
| Other CCs | 29 (20.3) | 42 (21.2) | 55 (18.2) | 16 (42.1) |
| Other Singletons | 4 (100) | 19 (100) | 22 (100) | 1 (100) |
| **Streptolancidin G** | | | | |
| **CC** | **Pre-PCV** | **Post-PCV** | **Carriage** | **IPD** |
| CC1146 | 48 (90.6) | 86 (100) | 121 (96.0) | 13 (100) |
| CC852 | 27 (100) | 51 (100) | 67 (100) | 11 (100) |
| CC5329 | 14 (93.3) | 23 (100) | 33 (97.1) | 4 (100) |

| | | | | |
|---|---|---|---|---|
| CC393 | 4 (100) | 5 (100) | 5 (100) | 4 (100) |
| CC5902 | 0 | 3 (2.2) | 3 (1.4) | 0 |
| CC5796 | 2 (15.4) | 0 | 2 (14.3) | 0 |
| CC14774 | 0 | 2 (11.8) | 2 (9.5) | 0 |
| CC909 | 1 (100) | 1 (100) | 2 (100) | 0 |
| Sing14823 | 1 (100) | 0 | 1 (100) | 0 |
| CC473 | 1 (25.0) | 0 | 1 (33.3) | 0 |
| Other CCs | 1 (50.0) | 1 (7.1) | 2 (15.4) | 0 |
| Other Singletons | 0 | 1 (100) | 1 (100) | 0 |
| **Streptolassin** | | | | |
| **CC** | **Pre-PCV** | **Post-PCV** | **Carriage** | **IPD** |
| CC289 | 64 (100) | 5 (100) | 3 (100) | 66 (100) |
| CC13854/15057 | 0 | 3 (100) | 3 (100) | 0 |
| CC404 | 1 (100) | 0 | 1 (100) | 0 |
| CC5936/14865 | 1 (100) | 0 | 1 (100) | 0 |
| Sing5359 | 1 (100) | 0 | 1 (100) | 0 |
| Sing14840 | 1 (100) | 0 | 1 (100) | 0 |
| CC5068 | 1 (33.3) | 0 | 0 | 1 (50.0) |

Note: The 10 most common clonal complexes in which the bacteriocin cluster was found are listed separately, and the remainder were pooled as 'Other'. Only bacteriocin clusters with significant differences in prevalence in Figure 2B and 2C are included in this table. IPD, invasive pneumococcal disease.

**Supplementary Table 10: Clonal complexes (CCs) and sequence types (STs) in the Icelandic dataset with multiple bacteriocin repertoires.**

| CC | Variable bacteriocins (CC) | Mixed STs | Variable bacteriocins (ST) |
|---|---|---|---|
| 236/271/320 | Streptococcin A, Streptococcin E | 271 | Streptococcin E |
| | | 1968 | Streptococcin A |
| 439 | Streptococcin A, Streptolancidin C, Streptolancidin D | 311 | Streptococcin A |
| | | 507 | Streptococcin A |
| | | 442 | Streptococcin A |
| | | 190 | Streptococcin A |
| 199 | Streptolancidin J, Streptosactin | 199 | Streptolancidin J, Streptosactin |
| 138/176 | Streptococcin A | 176 | Streptococcin A |
| 180 | Streptococcin A | 180 | Streptococcin A |
| 62 | Streptococcin A, Streptolancidin J | 62 | Streptolancidin J |
| 97 | Streptococcin A, Streptolancidin F, Streptolancidin J | 1635 | Streptolancidin J |
| 490 | Streptolancidin J | 2221 | Streptolancidin J |

| | | | |
|---|---|---|---|
| 124 | Streptococcin A | 124 | Streptococcin A |
| 433 | Streptocyclicin, Streptolancidin F, Streptolassin | 433 | Streptolancidin F |
| 30 | Streptocyclicin, Streptolancidin C, Streptolancidin D, Streptolancidin E, Streptolancidin G | 30 | Streptolancidin E |
| 392 | Streptocyclicin | 440 | Streptocyclicin |
| 156/162 | Streptocyclicin, Streptolancidin J | 162 | Streptolancidin J |
| 344 | Streptocyclicin, Streptolancidin F, Streptolancidin K | 10371 | Streptocyclicin, Streptolancidin F |
| | | 344 | Streptolancidin F, Streptolancidin K |
| 15 | Streptolancidin G | None | NA |
| 1262 | Streptolancidin J | 1262 | Streptolancidin J |
| 193 | Streptolancidin J | 1877 | Streptolancidin J |
| 113 | Streptococcin A, Streptolancidin F | 113 | Streptococcin A, Streptolancidin F |
| 113 | Streptococcin A, Streptolancidin F | 110 | Streptococcin A |
| 393 | Streptococcin A | None | NA |
| 66 | Streptolancidin G | None | NA |
| 315 | Streptolancidin G, Streptolancidin J | 386 | Streptolancidin J |
| 6524 | Streptolancidin J | 6524 | Streptolancidin J |
| 338 | Streptolancidin A, Streptolancidin D | None | NA |
| 63 | Streptolancidin D | None | NA |
| 432 | Streptolancidin G | 432 | Streptolancidin G |
| 205 | Streptolancidin J | 205 | Streptolancidin J |
| 306 | Streptococcin A | None | NA |
| 230 | Streptolancidin J | None | NA |
| 473 | Streptolancidin G | None | NA |

**Supplementary Table 11: Clonal complexes (CCs) and sequence types (STs) in the Kenyan dataset with multiple bacteriocin repertoires.**

| CC | Variable bacteriocins (CC) | Mixed STs | Variable bacteriocins (ST) |
|---|---|---|---|
| 5902 | Streptococcin A, Streptocyclicin, Streptolancidin B, Streptolancidin D, Streptolancidin E, Streptolancidin G, Streptolancidin J | 5902 | Streptocyclicin, Streptolancidin J |
| | | 5370 | Streptococcin A, Streptolancidin E |
| | | 840 | Streptolancidin D |
| | | 2052 | Streptolancidin E |
| | | 15056 | Streptolancidin G |
| 217 | Streptococcin A, Streptolancidin E | 613 | Streptococcin A, Streptolancidin E |
| 701 | Streptococcin A, Streptolancidin D, Streptolancidin J | 701 | Streptococcin A, Streptolancidin D |
| | | 5340 | Streptococcin A |
| 5339 | Streptococcin A, Streptocyclicin, Streptolancidin C, Streptolancidin D, Streptolancidin J | 5339 | Streptococcin A, Streptolancidin D |
| | | 844 | Streptococcin A, Streptocyclicin, Streptolancidin C, Streptolancidin D |
| | | 5367 | Streptolancidin J |
| | | 5268 | Streptococcin A, Streptolancidin C |
| 1146 | Streptolancidin E, Streptolancidin G, Streptolancidin J | 5952 | Streptolancidin E |
| | | 5396 | Streptolancidin G |
| 138/176 | Streptococcin A, Streptocyclicin, Streptolancidin A, Streptolancidin E, Streptolancidin J | 848 | Streptococcin A, Streptolancidin E, Streptolancidin J |
| 156/162 | Streptolancidin E, Streptolancidin K | 847 | Streptolancidin E, Streptolancidin K |
| 230 | Streptocyclicin, Streptolancidin D, Streptolancidin E, Streptolancidin F, Streptolancidin J | 230 | Streptolancidin D, Streptolancidin J |
| | | 700 | Streptocyclicin |
| | | 4351 | Streptolancidin E |
| 852 | Streptococcin A | 852 | Streptococcin A |
| 5258 | Streptococcin A | 5258 | Streptococcin A |
| 63 | Streptolancidin C, Streptolancidin J | 842 | Streptolancidin J |

| | | | |
|---|---|---|---|
| | | 2716 | Streptolancidin C |
| 347 | Streptococcin A, Streptocyclicin, Streptolancidin B, Streptolancidin C, Streptolancidin J | 6088 | Streptococcin A, Streptolancidin J |
| | | 2715 | Streptolancidin B, Streptolancidin J |
| | | 5769 | Streptococcin A, Streptolancidin J |
| | | 6095 | Streptolancidin J |
| | | 14817 | Streptococcin A, Streptolancidin J |
| 914 | Streptolancidin B | None | NA |
| 7053 | Streptococcin A, Streptolancidin C, Streptolancidin J | 5368 | Streptolancidin J |
| 702 | Streptolancidin B, Streptolancidin C | 702 | Streptolancidin B, Streptolancidin C |
| 854 | Streptococcin A, Streptocyclicin, Streptolancidin J | 854 | Streptococcin A, Streptocyclicin, Streptolancidin J |
| 499 | Streptolancidin C, Streptolancidin J | 499 | Streptolancidin C |
| | | 5907 | Streptolancidin J |
| 5329 | Streptocyclicin, Streptolancidin C, Streptolancidin G, Streptolancidin J, Streptolancidin K | 5329 | Streptolancidin C, Streptolancidin J |
| 338 | Streptolancidin D, Streptolancidin E, Streptolancidin J | 172 | Streptolancidin E |
| | | 2054 | Streptolancidin D, Streptolancidin J |
| 3460 | Streptococcin A, Streptocyclicin, Streptolancidin B | 14886 | Streptococcin A, Streptocyclicin |
| | | 3460 | Streptococcin A |
| 385 | Streptococcin A, Streptolancidin B, Streptolancidin J | 6097 | Streptolancidin B |
| | | 2713 | Streptococcin A |
| | | 3207 | Streptolancidin J |
| 989 | Streptolancidin D | 989 | Streptolancidin D |
| 14774 | Streptolancidin B, Streptolancidin G, Streptolancidin J | 6092 | Streptolancidin B, Streptolancidin G |
| | | 14774 | Streptolancidin J |
| 14892 | Streptococcin A, Streptolancidin C, Streptolancidin D, Streptolancidin E | None | NA |

| | | | |
|---|---|---|---|
| 2386 | Streptolancidin C, Streptolancidin D | 5331 | Streptolancidin D |
| 14858 | Streptocyclicin, Streptolancidin E | 14858 | Streptolancidin E |
| | | 14910 | Streptocyclicin |
| 4894 | Streptococcin A | 4894 | Streptococcin A |
| 5938 | Streptolancidin D | None | NA |
| 14930/15024 | Streptocyclicin, Streptolancidin C, Streptolancidin J | 14930 | Streptocyclicin, Streptolancidin C, Streptolancidin J |
| 5349 | Streptococcin A | None | NA |
| 5294 | Streptococcin A, Streptocyclicin | 5294 | Streptococcin A, Streptocyclicin |
| 703 | Streptocyclicin, Streptolancidin C, Streptolancidin J | 703 | Streptolancidin C, Streptolancidin J |
| Sing5373 | Streptolancidin C | 5373 | Streptolancidin C |
| 5372/15025 | Streptolancidin G | 5372 | Streptolancidin G |
| Sing14868 | Streptococcin A | 14868 | Streptococcin A |
| 5796 | Streptococcin A, Streptolancidin G, Streptolancidin J | 5796 | Streptococcin A |
| 5560/6090/6103 | Streptococcin A | 6103 | Streptococcin A |
| 193 | Streptolancidin C, Streptolancidin J | None | NA |
| 1766 | Streptococcin A | None | NA |
| 1264 | Streptolancidin D | 1264 | Streptolancidin D |
| 3735 | Streptocyclicin | None | NA |
| 14846/14876 | Streptococcin A, Streptolancidin D | 14846 | Streptococcin A, Streptolancidin D |
| 5798/5879 | Streptococcin A, Streptolancidin B | 5798 | Streptococcin A, Streptolancidin B |
| 5321/14966 | Streptolancidin C | None | NA |
| 393 | Streptolancidin J | None | NA |
| 547 | Streptolancidin D, Streptolancidin J | None | NA |
| 3518 | Streptococcin A, Streptolancidin B | None | NA |
| 3983 | Streptococcin A | 3983 | Streptococcin A |
| 5266 | Streptolancidin J | None | NA |
| 5901 | Streptococcin A | 14976 | Streptococcin A |
| 5398/14814 | Streptolancidin C | None | NA |
| 473 | Streptolancidin G | None | NA |

| | | | |
|---|---|---|---|
| 5839/14990 | Streptolancidin J | None | NA |
| 5068 | Streptolancidin J, Streptolassin | None | NA |
| Sing5376 | Streptolancidin J | 5376 | Streptolancidin J |
| 849/5343/5351 | Streptococcin A | None | NA |

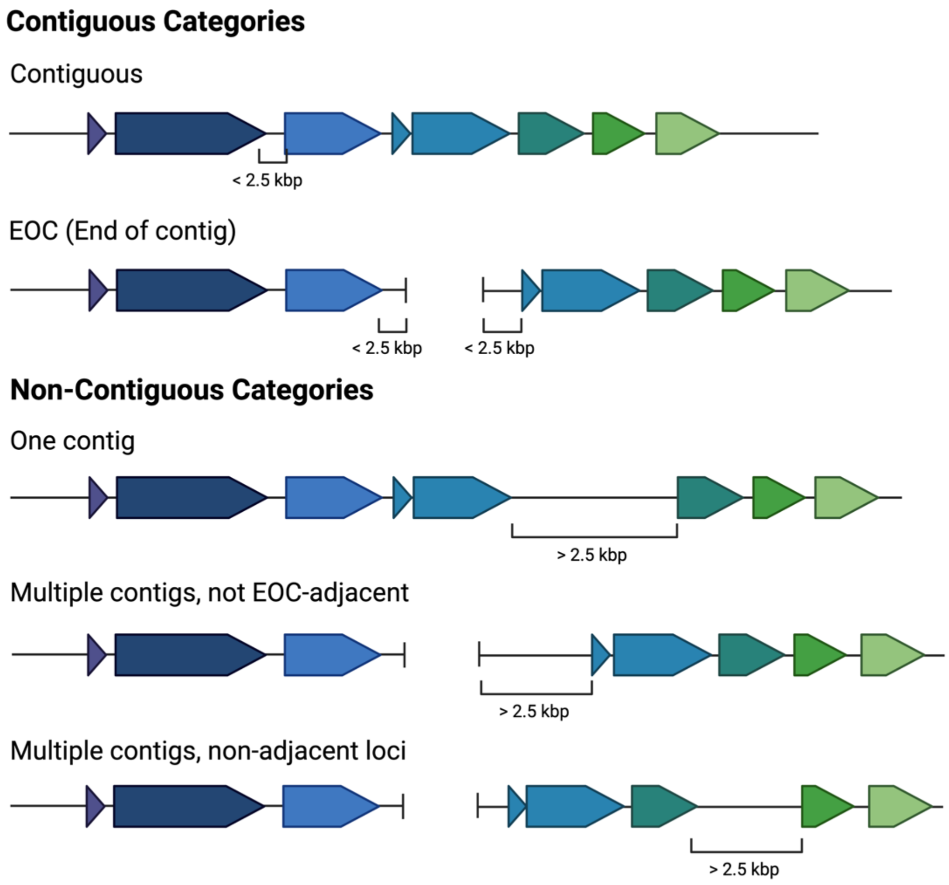


**Supplementary Figure 1:** Illustration of a hypothetical bacteriocin biosynthetic gene cluster and the various cluster contiguity categories quantified in Supplementary Table 6 (figure created using BioRender).
